# Genome assembly of the dyeing poison frog provides insights into the dynamics of transposable element and genome-size evolution

**DOI:** 10.1101/2023.11.06.565769

**Authors:** Carolin Dittrich, Franz Hölzl, Steve Smith, Chloe A. Fouilloux, Darren J. Parker, Lauren A. O’Connell, Lucy S. Knowles, Margaret Hughes, Ade Fewings, Rhys Morgan, Bibiana Rojas, Aaron A. Comeault

## Abstract

Genome size varies greatly across the tree of life and transposable elements are an important contributor to this variation. Among vertebrates, amphibians display the greatest variation in genome size, making them ideal models to explore the causes and consequences of genome size variation. However, high-quality genome assemblies for amphibians have, until recently, been rare. Here, we generate a high-quality genome assembly for the dyeing poison frog, *Dendrobates tinctorius*. We compare this assembly to publicly-available frog genomes and find evidence for both large-scale conserved synteny and widespread rearrangements between frog lineages. Comparing conserved orthologs annotated in these genomes revealed a strong correlation between genome size and gene size. To explore the cause of gene-size variation, we quantified the location of transposable elements relative to gene features and find that the accumulation of transposable elements in introns has played an important role in the evolution of gene size in *D. tinctorius*, while estimates of insertion times suggest that many insertion events are recent and species-specific. Finally, we show that the diversity and abundance of transposable elements in poison frog genomes can complicate genotyping efforts that rely on repetitive elements as sequence anchors. Our results show that transposable elements have clearly played an important role in the evolution of large genome size in *D. tinctorius*. Future studies are needed to fully understand the dynamics of transposable element evolution and to optimise primer or bait design for cost-effective population-level genotyping in species with large, repetitive genomes.

**Significance:** Amphibians display more variation in genome size than any other vertebrate lineage. Complexities associated with large genomes frequently hamper genome assembly and population genetic studies. Here we use long-read HiFi sequences to generate a high-quality 6.3 Gbp genome assembly of the poison frog *Dendrobates tinctorius*. We use this genome and leverage comparative genomics and *de novo* annotations to quantify aspects of genome evolution driven by repetitive transposable genetic elements. Our results provide support for the dynamic role that transposable elements play in driving the evolution of “genomic gigantism” in amphibians. We also show how transposable elements can be leveraged for cost-efficient population genetic studies using limited input material.

## Introduction

Interspecific variation in genome size is a fundamental feature of biodiversity, and transposable elements play an important role in contributing to variation in genome size and structure (Kidwell 2002; Hawkins et al. 2006; Lee and Kim 2014). While frequently and historically referred to as “junk” DNA, it has been long known that the evolution of transposable elements can have profound effects on an organism’s phenotype. For example, the ability of transposable elements to drive gene expression and mosaic coloration in maize accompanied their discovery in the late 1940s by Barbara McClintock (McClintock 1950). With the advancement of computational capabilities and the availability of cost-effective genetic methods, evidence of the phenotypic effects of transposable elements has increased. It is now widely appreciated that transposable elements can have a profound effect on evolution via their ability to underlie the evolution of exon structure, telomeres, gene expression, and ultimately, adaptation or speciation (Almojil et al. 2021; Casacuberta and González 2013; Feschotte 2008; Serrato-Capuchina and Matute 2018). The activity of transposable elements has even recently been linked to plasticity and adaptation to global change (Pimpinelli and Piacentini 2020; Rey et al. 2016). For a comprehensive classification and in-depth review of the impact transposable elements can have on the genome, we recommend the recent works of (Almojil et al. 2021; Bourge et al. 2018; Kidwell and Lisch 1997; Sotero-Caio et al. 2017; Wicker et al. 2007).

Characterising the genomic landscape of transposable elements—such as the abundance of different types of elements and where they are located in the genome relative to genes, exons, and introns—is one approach that can be used to illuminate aspects of their evolution. With the increasing availability of whole-genome sequence data collected from many species, studies are increasingly quantifying the genomic landscape and “ecology” of transposable elements (Lamichhaney et al. 2021; Stitzer et al. 2021; Gozashti et al. 2023). A general pattern emerging from these studies is that different types of transposable elements contribute to genome evolution in different species. For example, long terminal repeat (LTR) retrotransposons play an important role in genome size variation among plants (Lee and Kim 2014); while long and short interspersed nuclear elements (LINES and SINEs, respectively) are more abundant than LTRs in mammals (Chalopin et al. 2015; Platt et al. 2018), and SINEs are nearly absent in amphibian genomes (Zuo et al. 2023). In addition to the types of repetitive elements present within genomes, estimates of the timing of when different repetitive elements insert themselves vary both across species (Sun et al. 2015) and among the types of repetitive elements present within single genomes (Sun et al. 2015; Stitzer et al. 2021). Finally, different types of repetitive elements can display different “ecologies”, with some being prone to insert themselves into non-random locations in the genome, such as in intergenic regions, promotors, or introns (Bourque et al. 2018; Stitzer et al. 2021). Gaining a better understanding of the “ecology” displayed by different repetitive elements across different species is important because it will help facilitate a predictive understanding of how repetitive elements contribute to the evolution of genomic and genetic variation.

Among vertebrates, amphibians exhibit remarkable variation in genome size, surpassing that of any other group (Liedtke et al. 2018). The variation in amphibian genome size is influenced by the activity and accumulation of transposable elements (Sotero-Caio et al. 2016). However, the limited availability of high-quality amphibian genome assemblies—in large part due to the challenges of assembling large repetitive genomes with short-read sequencing technologies—makes the evolutionary dynamics and genomic ecology of TEs, alongside their impact on phenotypes and adaptation, difficult to test (however, see Zuo et al. 2023).

As complete genomes are often lacking for amphibians, the research community were early adopters of reduced representation library sequencing for studies of species delimitation and population genetics (Dufresnes et al. 2018; Homola et al. 2019; Nunziata and Weisrock 2018). Given their transposable element-rich genomes, it is surprising that methods that leverage repetitive elements, such as MobiSeq (Rey-Iglesia et al. 2019), have not yet been applied to amphibians. MobiSeq is a method for constructing a reduced representation library by targeting the flanking regions of transposable elements to identify single nucleotide polymorphisms (SNPs) and genotypes (Rey-Iglesia et al. 2019). This approach requires minimal DNA input and does not require a reference genome, making it useful for investigating a wide array of research questions with regards to population genetics, evolutionary dynamics and ecological interactions.

Poison frogs (Family Dendrobatidae) are a group of Central- and South American forest-dwelling amphibians with complex social behaviour and elaborate parental care (Stynoski et al. 2015). A thriving research community have focused on this model clade for studying the effects of natural selection on phenotype, particularly the bright colouration coupled with chemical defences that protects some species from predation (Noonan and Comeault 2009; Chouteau et al. 2011; Maan and Cummings 2012; Lawrence and Rojas et al. 2019). A solid natural history background stemming from field observations done in the late 1900s (e.g. Sexton 1960, Silverstone 1973; Wells 1980; Myers and Daly 1983; Donnelly 1989; Summers 1989) had laid a firm foundation to ask both ultimate and proximate research questions. For example, research on the ultimate factors shaping their communication and territorial behaviour, parental care, and space use, as well as the behaviour of their larvae, has been steadily gaining traction (e.g., Pröhl 2005; Summers et al. 2006; Amézquita et al. 2011; Ringler et al. 2013; Schulte et al. 2013; Tumulty et al. 2014; Rojas 2014; Stynoski et al. 2014; Carvajal-Castro et al. 2021; Fouilloux et al. 2021). More recently, studies on the proximate mechanisms of such behaviours, i.e., neurobiology of egg provisioning, tadpole transport and tadpole aggression (Fischer and O’Connell 2020; Fischer et al. 2019, 2020), and hormonal correlates of care, territoriality and space use (Fischer and O’Connell 2020; Pašukonis et al. 2022; Rodríguez et al. 2022), now provide a more holistic understanding of what makes these frogs unique. However, the lack of genomic resources has precluded some of these topics from being addressed in depth. The one genome available for dendrobatid frogs (*Oophaga pumilio*) is highly fragmented, but still shows that transposable elements comprise a large portion of it (Rogers et al. 2018). Having access to reference genomes from more poison frogs would allow for more comprehensive approaches to questions related to demography, conservation, behavioural ecology, disease dynamics and adaptation (Brandies et al. 2019).

In this study, we generate a high-quality reference genome for the dyeing poison frog, *Dendrobates tinctorius*. This species is aposematic—with drastic differences in coloration and toxicity across populations (Lawrence and Rojas et al. 2019)—and displays complex social behaviours typical of many poison frog species including male parental care and territoriality (Rojas and Pašukonis 2019; Fouilloux et al. 2021; Rojas 2014; 2015). Here, we first leverage our genome assembly and three publicly available chromosome-scale assemblies of species belonging to Hyloidea to provide evidence for both large regions of synteny alongside significant structural evolution. This analysis also revealed that the evolution of genome size across these species is correlated with gene size, with *D. tinctorius* having both the largest genome and the longest genes. We then annotate transposable elements in the *D. tinctorius* genome to explore the genomic landscape of their evolution and find that transposable elements are more abundant in introns than in exons, likely contributing to the evolution of large genes in *D. tinctorius*. Finally, we explored the usefulness of the *D. tinctorius* reference genome in leveraging population genetic information from cost-effective mobile element sequencing (MobiSeq, Rey-Iglesia et al. 2019). Applying this method to tadpoles collected from a wild population revealed that the highly repetitive nature of the *D. tinctorius* genome can generate challenges to MobiSeq genotyping, with a high degree of missing data across sites and individuals. We hope that the ability to use the *D. tinctorius* genome as a resource will refine the development of markers—such as additional mobile-element tags or alternate approaches such as baited capture—for genotyping individuals, thereby broadening the scope of behavioural ecology and population genomic research in amphibians.

## Results

### Genome assembly, quality control

Our final *Dendrobates tinctorius* assembly consists of 6.356 Gb assembled into 830 contigs. This assembly has a contig N50 and L50 of 32.539 Mb and 56 contigs, respectively, with a maximum contig size of 131.4 Mb. Our assembly is also highly accurate and complete, achieving an error rate of less than 0.0001—as indicated by a QV score of 41—and containing 4,796 of 5,310 curated tetrapod BUSCO genes (BUSCO summary: C:90.3% [S:88.4%, D:1.9%], F:2.8%, M:6.9%, n:5310). While our assembly does not contain complete chromosomes, the contiguity and BUSCO scores were comparable to chromosome-level assemblies of other anurans available on NCBI (fig. 1).

**Fig. 1.**
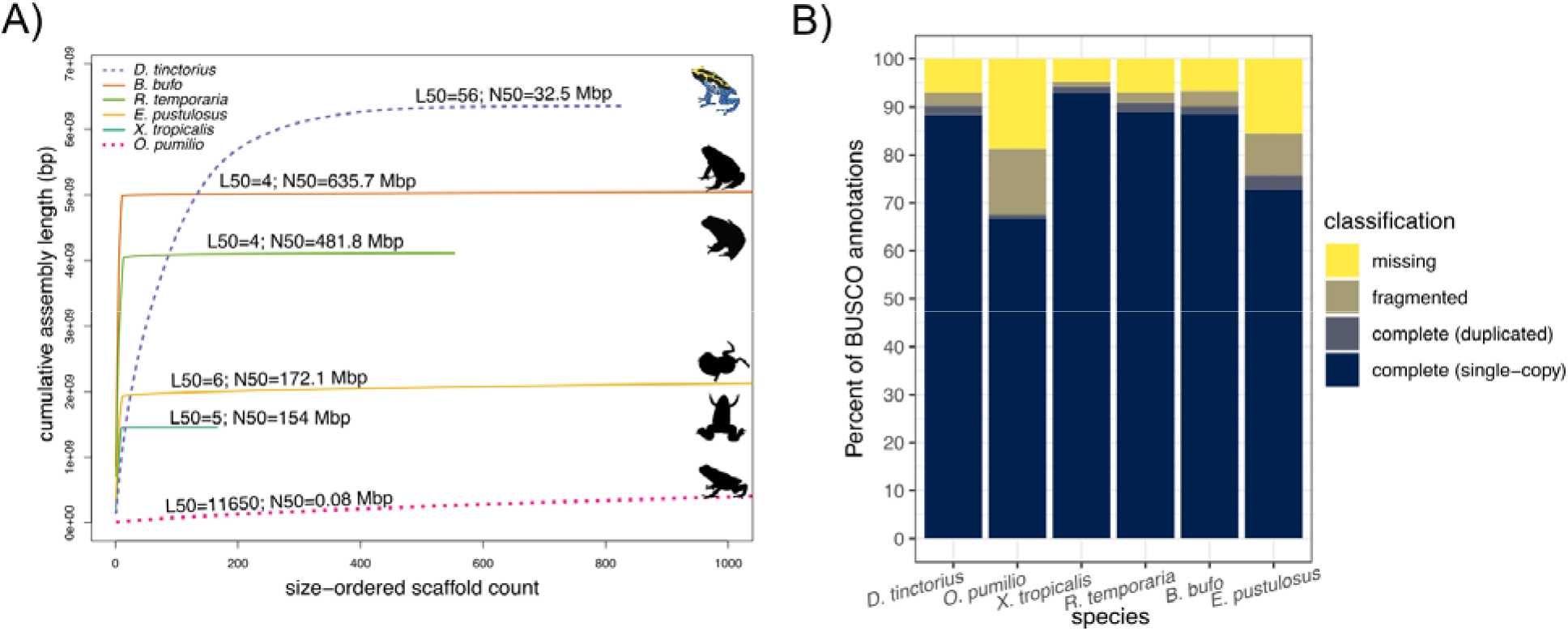
*Dendrobates tinctorius* assembly completeness. (**A**) Cumulative assembly size in relation to size-ordered scaffolds for *D. tinctorius* and other published anuran genomes. (**B**) BUSCO “tetrapoda” gene set completeness for the same genomes summarised in (A).

### Synteny between D. tinctorius and other Hyloidea genomes

To explore patterns of conserved synteny between our assembly and closely related Anuran species, we compared the *D. tinctorius* assembly to chromosome-scale assemblies of *Bufo gargarizans* (asiatic toad; Bufonidae), *Eleutherodactylus coqui* (common coquí; Eleutherodactylidae), and *Engystomops pustulosus* (túngara frog; Leptodactylidae). *Bufo gargarizans*, *El. coqui*, and *En. pustulosus* shared a common ancestor with *D. tinctorius* ∼65 Mya (Feng et al. 2017), and, while these four species have significantly diverged from one another, the former three were used for comparisons because they represent the three most closely related genera to *Dendrobates* that currently have publicly available chromosome-scale genome assemblies. Orthology-guided synteny analyses based on the tetrapoda BUSCO geneset in GENESPACE provided evidence of both broad-scale synteny and genome evolution between the *D. tinctorius* and the three other species we analysed (Fig. 2). The number of synteny blocks identified between *D. tinctorius* and the chromosome-scale assemblies ranged from 116 (*En. pustulosus*) to 164 (*B. gargarizans*), with an average size of each block of 27.36 Mb (*En. pustulosus*) to 17.86 Mb (*B. gargarizans*). We focused this analysis on scaffolds of the *D. tinctorius* assembly that were greater than 20 Mb in length (see Methods), and these *D. tinctorius* scaffolds have median and mean lengths of 32.64 Mb and 41.81 Mb, respectively. Synteny between *D. tinctorius* scaffolds and the chromosome-scale assemblies therefore tends to span roughly half of entire *D. tinctorius* scaffolds, on average, while in many cases entire scaffolds showed collinearity with chromosomal regions of the chromosome-scale assemblies (fig. 2B and C). In addition to collinear regions, we observed numerous rearrangements between *D. tinctorius* scaffolds and chromosomes of the other Anuran genomes we analysed: for example, Figure 2B highlights rearrangements between *D. tinctorius* scaffolds and *B. gargarizans*’s chromosome 1 (inversions highlighted in salmon pink colour). Finally, we observed a single instance where a *D. tinctorius* scaffold contained synteny blocks that mapped to different chromosomes of a chromosome-scale assembly (*El. coqui* chromosomes 1 and 4; fig. 2C). These results suggest that—at least at the course scale of ∼65 My of evolution— rearrangements within chromosomes are more numerous than changes in the overall evolution of chromosome number via fission and/or fusion events.

**Fig. 2.**
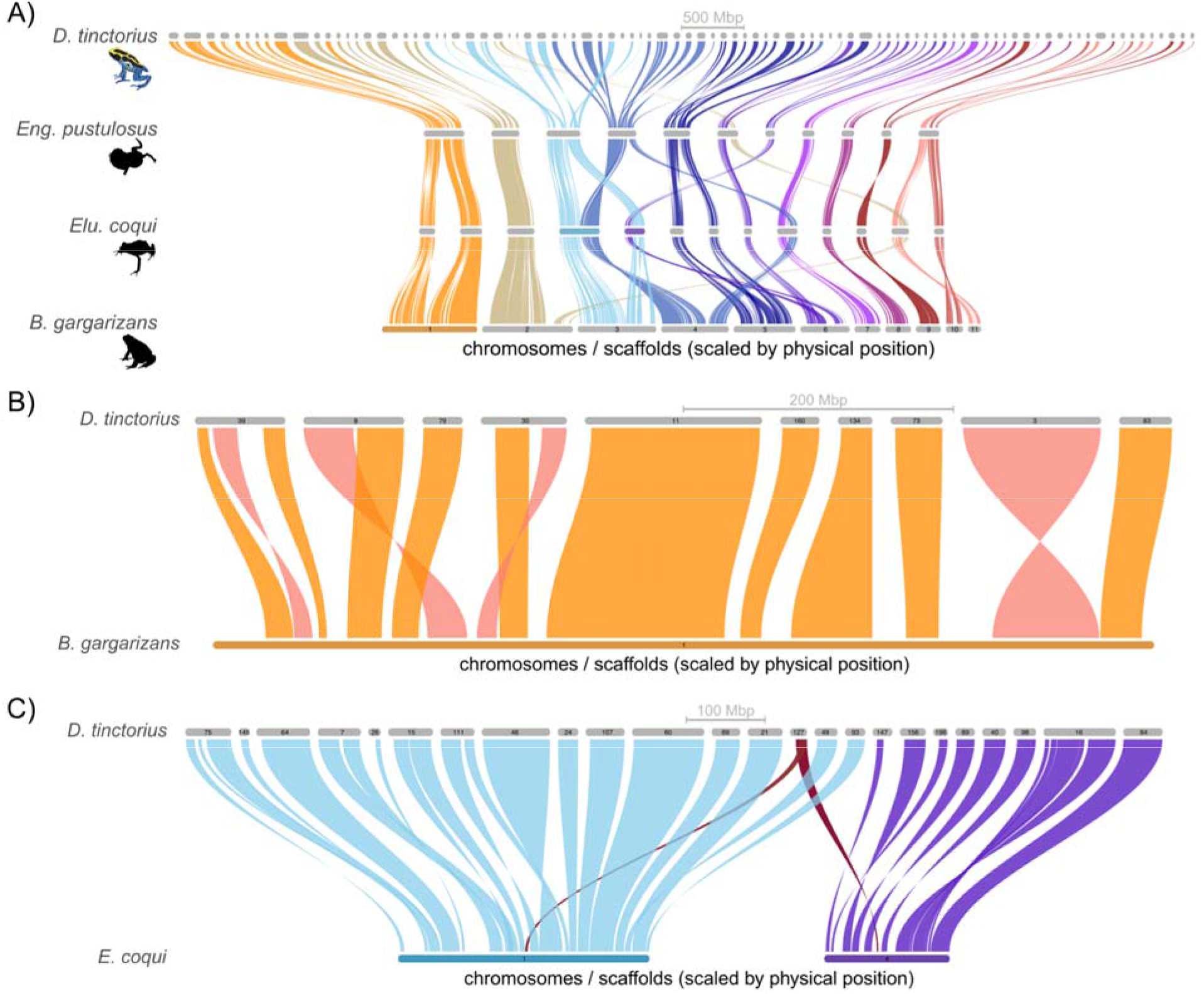
Riparian synteny plots generated using BUSCO gene regions as anchors showing broad patterns of synteny and rearrangements between *D. tinctorius* scaffolds and three publicly available chromosome-scale frog assemblies (**A**). (**B**) Example *B. gargarizans* chromosome showing both synteny and structural evolution with *D. tinctorius* scaffolds. Inversions in panel B are shown in salmon pink, collinear regions in orange. (**C**) Insertion of regions on *D. tinctorius* scaffold 127 (highlighted in red) into chromosomes 1 and 4 of the *E. coqui* assembly.

Our *D. tinctorius* assembly is larger than any of the four chromosome-scale genomes we used in the analyses presented above (*B. gargarizans*, *El. coqui*, *En. pustulosus*, + *Xenopus tropicalis*), with genome size ranging from 1.451 Gb (*X. tropicalis*) to 6.356 Gb (*D. tinctorius*). We therefore explored how genome size evolution affects the size of gene-regions by comparing the size of BUSCO annotations in each of these genomes. Across 2,035 single cop orthologs that were present in all genomes, *D. tinctorius* orthologs were 99.5 to 205.9% longer, on average, than the same orthologs in the *B. gargarizans* and *X. tropicalis* assemblies. This percent difference in gene size was negatively correlated with genome size (Spearman’s rank correlation test; *P* = 0.083), with larger genomes (i.e., *B. gargarizans* versus *D. tinctorius*) showing less difference in genes size than when genomes differed in size (i.e., *X. tropicalis* versus *D. tinctorius*). In general, we found a positive correlation between genome size and the average size of BUSCO gene regions (rho = 0.99; *P* = 0.017, fig. 3), indicating that the evolution of larger genomes in *D. tinctorius* and *B. gargarizans* has resulted in the evolution of larger gene regions.

**Fig. 3.**
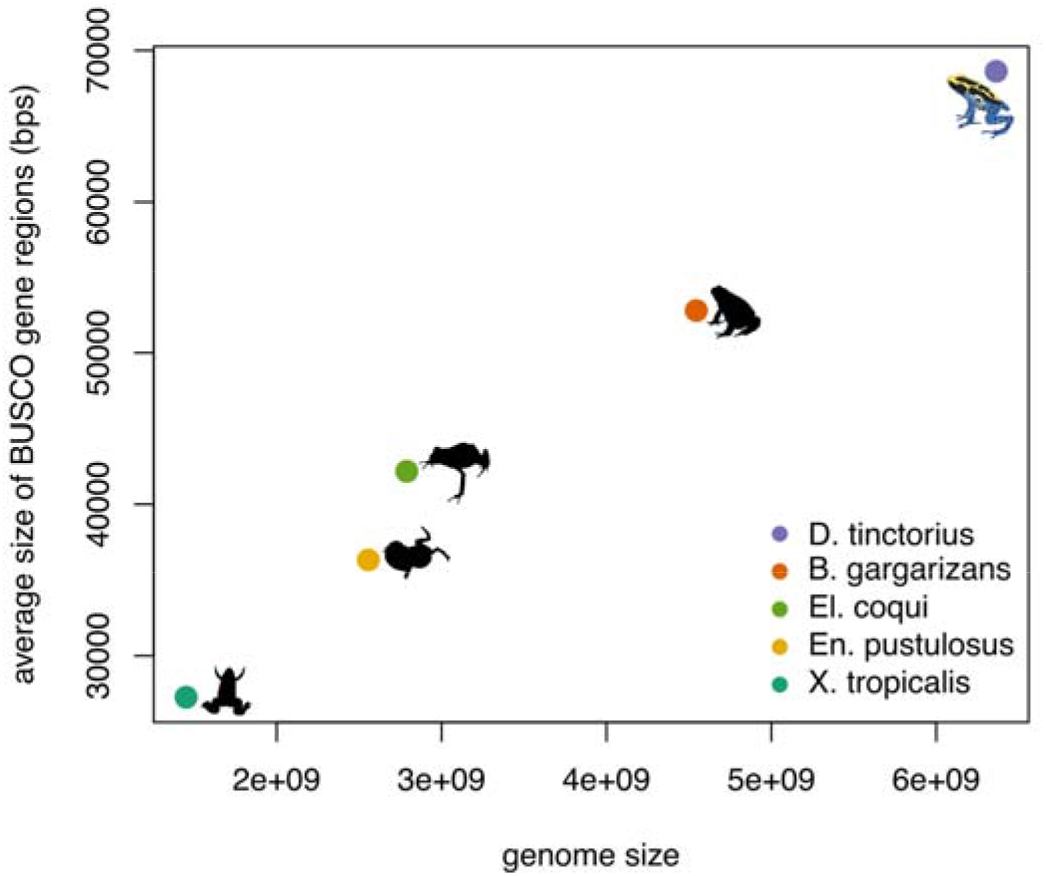
Scatterplot illustrating the relationship between genome size and the average size of annotated BUSCO gene regions in the *D. tinctorius* assembly and four publicly available chromosome-scale frog assemblies. *Dendrobates tinctorius* has both the longest genome and the longest average BUSCO gene length.

### Genome annotation and transposable element diversity

The majority of the *D. tinctorius* genome consists of repetitive elements: annotation with RepeatModeler + RepeatMasker identified 74.72% of our assembly as repetitive. Many of the repeats classified by RepeatMasker could not be assigned to specific types of repetitive elements (39.15%). Among assigned elements, 23.13% were retroelements and 12.45% were DNA transposons. The most common retroelements classified by RepeatMasker were LTR retrotransposons (15.69% of the assembly) and LINEs (7.4%). LTR retrotransposons *Gypsy*/*DIRS1* were the most abundant superfamily (13.79% of the assembly), while DNA transposons made up 12.45% of the assembly. Of DNA transposons, *Tc1-IS630-Pogo* (*Tc1*) and *hobo-Activator* (*hAT*) elements were the most abundant (6.57% and 3.91% of the assembly, respectively).

Independent analyses with LtrDetector annotated 262,486 LTRs spanning 2,343,246,106 bp, or 36.9% of our assembly. While this represents a higher percentage of LTRs in our assembly than when annotated with RepeatMasker, consistent with RepeatMasker, the most abundant LTR elements were found to belong to the *Gypsy*/*DIRS1* superfamily (99,884 elements, 819.4 Mbp total), followed by *BEL*/*Pao* elements (1,762 elements, 13.9 Mbp total). Also consistent with results from RepeatMasker, 66,046 of the LTR elements annotated by LtrDetector (spanning 588 Mbp) had BLAST matches to the “Unknown” category of elements identified by RepeatModeler.

To better understand the role that repetitive elements have played in the evolution of *D. tinctorius* genome structure, we compared the location of repetitive elements relative to *de novo* gene annotations we generated using the BRAKER2 pipeline. Of the 11,331,718 repeat annotations generated by RepeatMasker, 115,330 (1.02%) overlapped with coding DNA sequence (CDS) and 4,872 of these had a reciprocal overlap of at least 75% (table 1 and table 2, respectively). More repetitive elements overlapped with introns compared to CDS (525,441 elements; ^2^ = 270,118; *P* < 2.2 x 10-16); however, proportionally fewer TEs that overlapped introns showed reciprocal overlaps of greater than 75% compared to those overlapping CDS (1.05 versus 4.22% respectively; ^2^ = 5,956.1; *P* < 2.2 x 10-16). These patterns are likely due to the fact that introns span nearly 4 times the number of bases and have a median size nearly twice that of CDS features (266 Mb versus 69 Mb and 113 versus 66 bp, respectively).

**Table 1.** Counts of repeat elements annotated using RepeatModeler + RepeatMasker that overlap with coding DNA sequence (CDS) and introns annotated in the *D. tinctorius* assembly.

| Region<br>of<br>overlap | All elements | LTRs | LINEs | *Tc1*<br>transposons | *hAT*<br>transposons |
| --- | --- | --- | --- | --- | --- |
| CDS | 115,330 | 3,500 | 1,826 | 10,227 | 507 |
| Introns | 528,441 | 42,844 | 27,152 | 45,806 | 2,907 |
| Total N<br>elements | 11,331,718 | 1,048,437 | 702,556 | 897,264 | 265,361 |

**Table 2.**
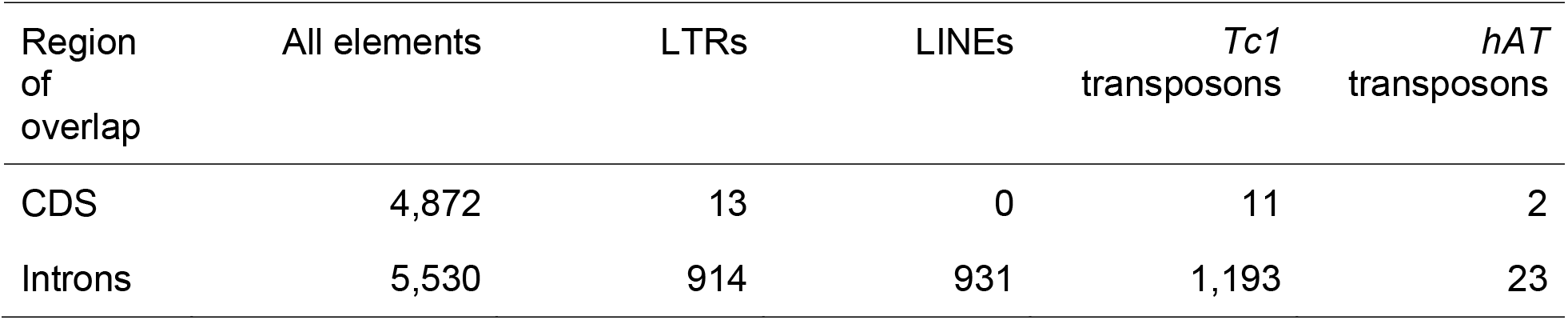
Counts of repetitive elements that overlap with CDS and introns (see Table 1) that show greater than 75% reciprocal overlap with those features.

| Region of overlap | All elements | LTRs | LINEs | *Tc1* transposons | *hAT* transposons |
| --- | --- | --- | --- | --- | --- |
| CDS | 4,872 | 13 | 0 | 11 | 2 |
| Introns | 5,530 | 914 | 931 | 1,193 | 23 |

We next considered positions of the four most abundant classes/families of repetitive elements annotated by RepeatMasker in the *D. tinctorius* genome—LTRs, LINEs, *Tc1* DNA transposons, and *hAT* DNA transposons—in relation to CDS and introns. Of these four classes/families of repeats, LTRs and *Tc1* transposons showed the highest overlap with CDS and introns, with 0.33% (3,500) of LTR elements and 1.14% (10,227) of *Tc1* transposons showing at least partial overlap with CDS, and 4.09% (42,844) and 5.11% (45,806) overlapping with introns, respectively. Only *Tc1* transposons were enriched in the proportion overlapping with CDS and introns compared to all non-*Tc1* elements (CDS: 1.14% versus 1.01%, respectively; introns: 5.27% versus 4.65%, respectively).

Given that a large portion of the *D. tinctorius* assembly is comprised of LTR elements— 15.69% to 36.9% of the assembly annotated by RepeatModeler or LtrDetector, respectively— we next explored length distributions of these elements and estimated the timing of their insertion by estimating divergence between the left and right LTR of each element, assuming a substitution rate of 2.5 x 10^-9^ substitutions per site per year (Lau et al. 2020). Average estimates of insertion times for different types of LTR retroelements ranged from 10 to 28 Mya (fig. 4). Retroelements with BLAST matches to the *DIRS* order of elements had the oldest average estimated insertion time (mean = 23.35 Mya, 95% empirical range = 1 - 60.4 Mya) while *ERV1* elements had the youngest insertion times (mean = 10.4 Mya, 95% empirical range = 0 - 45.1 Mya). The broad range of insertion times we estimated indicate that some LTR retroelements are old and may be shared with other species of poison frog, while others are young, potentially active, and species-specific.

**Fig. 4.**
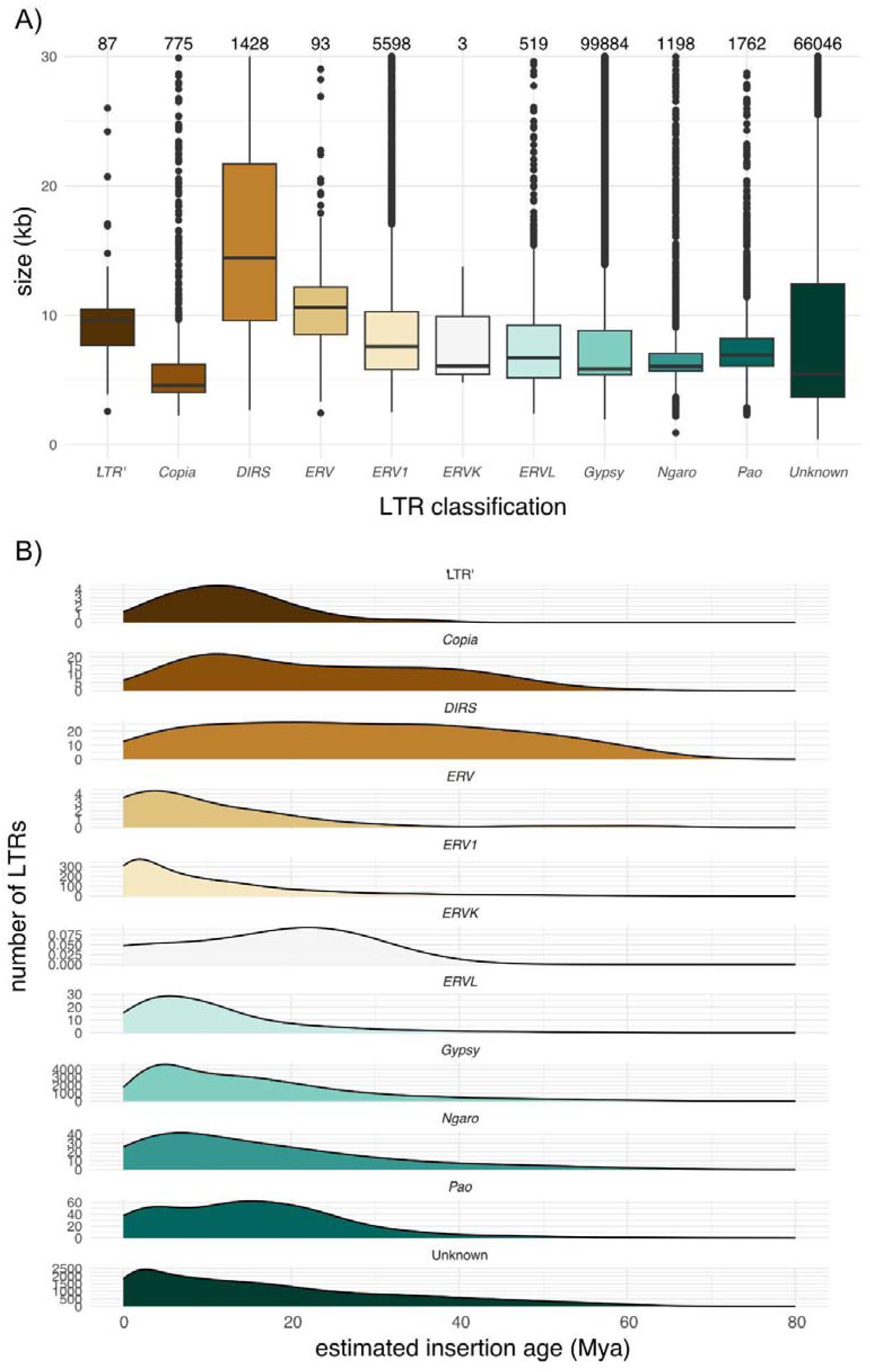
Long terminal repeat (LTR) retroelement lengths **(A)** and insertion times **(B)**. LTRs were annotated with LtrDetector and insertion times were estimated from divergence between the left and right LTR regions of each element independently assuming a substitution rate of 2.5 x 10^-9^ substitutions per site per year. In (A) the total number of elements classified to each type is given above each boxplot.

### A test of the utility of MobiSeq for population genetics

We next used the *D. tinctorius* reference genome to compare the genotyping success of three genotyping approaches using a MobiSeq dataset generated from 87 tadpoles collected from 17 phytotelmata at the Nouragues Research Station, French Guiana. Specifically, we compared the number of usable SNPs generated by either (1) de novo assembly of MobiSeq reads, (2) mapping the reads to the other poison frog genome currently available on NCBI (*O. pumilio*), or (3) mapping the reads to the *D. tinctorius* genome we generated as part of this study. We also called SNPs using two approaches: either the stacks pipeline (Catchen et al. 2011, 2013) designed to assemble RADseq data de novo or using a reference genome; or the original MobiSeq pipeline using the program “analysis in next generation sequencing data’’ (ANGSD) mapping sequence reads to the *D. tinctorius* assembly (Rey-Iglesia et al. 2019) (details in Methods).

The number of SNPs differed considerably between the two primers and different mapping approaches (de novo vs. reference genome), and showed high degrees of missing genotypes when genotyped using the stacks pipeline (LINE109: 93.21 ± 3.06%, TE644: 90.46 ± 1.46%; mean percent missing data across sites ± sd) and low coverage (supplementary table S1, Supplementary Material online). Regions amplified with the LINE109 primer resulted in considerably fewer SNPs than the TE644 primer. When we mapped sequences to the *O. pumilio* genome, gstacks only incorporated 10-12% of the reads and called considerably fewer SNPs than either the de novo approach or mapping to the *D. tinctorius* genome. The de novo approach with 50bp fragments called the highest numbers of SNPs (n SNPs TE644 = 92,433); in contrast, the 100bp fragments called the lowest numbers (n SNPs TE644 = 15,987). These differences were expected due to the large loss of SNPs due to truncation. Mapping to the *D. tinctorius* reference genome for both primers called the second most SNPs (n SNPs; LINE109 = 2,476, TE644 = 82,863), but missingness was still high and coverage low (supplementary table S1 and S2, Supplementary Material online). After applying a strong filter, where SNP had to be present in at least 50% of individuals, the numbers of SNP dropped dramatically (table S2 and fig. 5B).

**Fig. 5.**
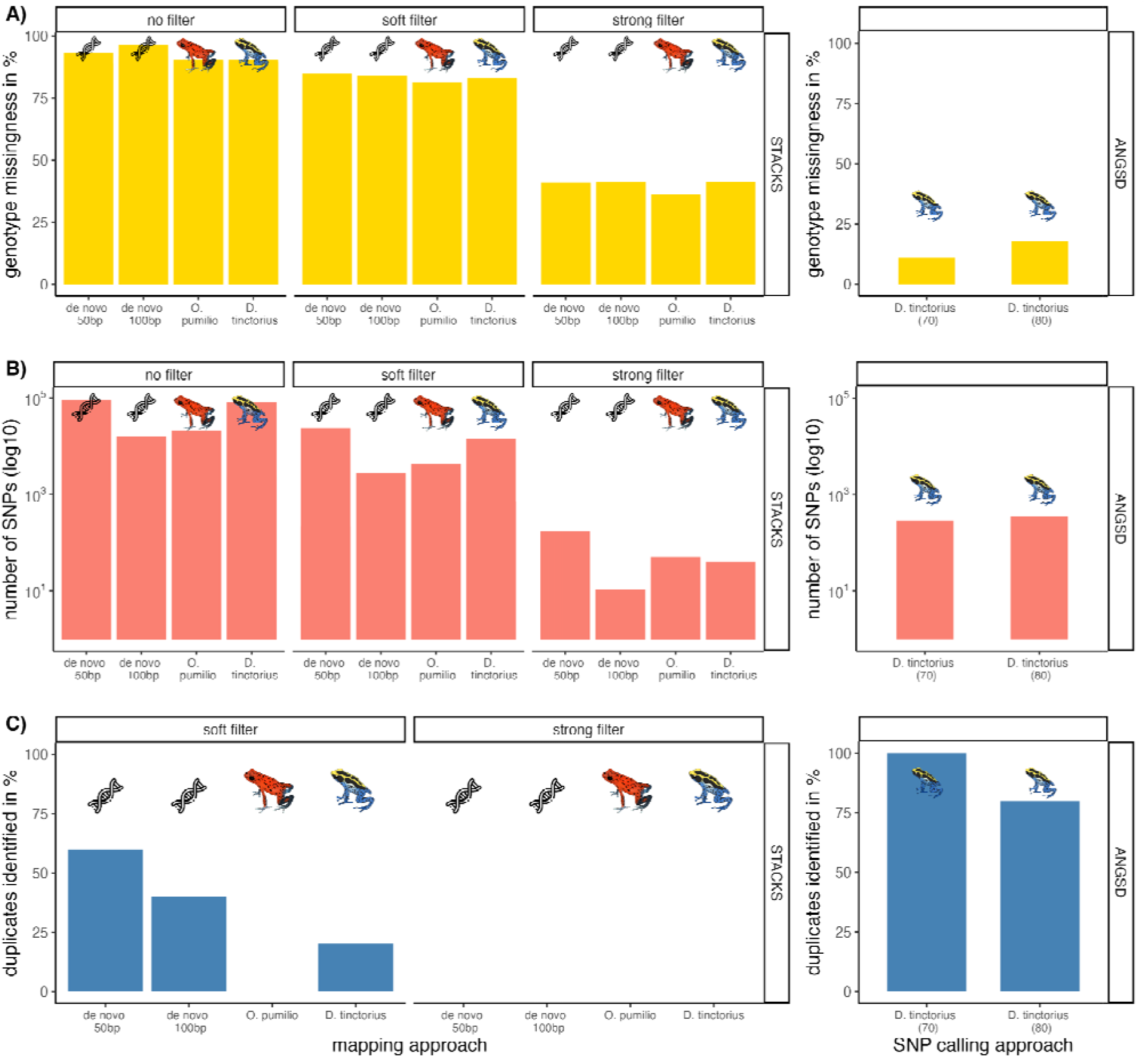
Summary of **(A)** genotype missingness in %, **(B)** number of SNPs (log scale to account for the large differences) and **(C)** percentage of duplicates found in the dataset using sequences derived from MobiSeq with the TE644 primer. Shown are the two pipeline approaches, stacks on the left (de novo assembly and mapping to reference genomes of *Oophaga pumilio* or *Dendrobates tinctorius*) with no filter SNPs, soft filter (filter_rad) and strong filter (vcftools); and ANGSD on the right (calling of SNPs present in 70 or 80 individuals). For details see Methods.

The SNP calling with the ANGSD pipeline gave 17.67 ± 0.17% missing genotypes and 345 SNPs were called when the output was restricted to be present in at least 70 individuals. If SNPs were restricted to be present in at least 80 individuals, 291 SNPs could be called with 11.07 ± 0.08% genotype missingness (fig. 5A and B).

We used our called SNP dataset to test their usefulness in parentage analysis. Specifically, we ran a relatedness analysis with COLONY (Jones and Wang 2010) on our dataset of 87 tadpoles. We included 5 sets of duplicate samples to test whether these samples will be reliably grouped together in our analysis. In general, we found significant differences in COLONY results depending on which SNP dataset we used for the analysis. Due to the low number of SNPs called with the LINE109 primers, we focused our COLONY analyses to data obtained using the TE644 primers. We considered the successful resolution of our duplicate samples, grouped together with a probability of 0.9 or greater, as evidence that a SNP data set was reliable for parentage analysis. Generally, reliability was low when using SNPs that were genotyped using Stacks: neither the de novo nor the reference genome mapping approaches reliably identified the 5 duplicates in the dataset with high probability (fig. 5C). When genotyped using the ANGSD pipeline, COLONY identified 4/5 duplicates when SNPs from at least 80 individuals were retained, and 5/5 duplicates when 70 individuals were the minimum threshold for calling SNPs, suggesting that the more complete datasets generated with ANGSD—with only 11-17% missing genotypes—provided sufficient information for identifying full-sibs (fig. 5A and C). This method also gave more realistic estimated population sizes than the stacks method (supplementary table S2, Supplementary Material online).

## Discussion

Access to genomic resources and tools holds the potential to transform our understanding of the ecology, evolution, life-history, and conservation of amphibians (e.g. Liedtke et al. 2018; Womack et al. 2019; Schloissnig et al. 2021; Kosch et al. 2022); yet amphibians have lagged behind other groups of animals in available genomic resources (Hotaling et al. 2021). The lack of genomic resources for frogs and toads (Anurans), in particular, is at least in part due to some species possessing large and complex genomes (Rogers et al. 2018; Nowoshilow et al. 2018; Sun et al. 2015): among frogs and toads genome size is highly variable ranging from 0.99 Gb in the plains spadefoot toad, *Spea bombifrons* (GenBank accession GCA_027358695.2) to 10.2 Gb in the mountain yellow-legged frog, *Rana muscosa* (GenBank accession GCA_029206835.1). Here we report a highly contiguous 6.8 Gb genome assembly of the dyeing poison frog, *D. tinctorius*. Using multiple approaches to annotation, we show that over three quarters of the *D. tinctorius* genome consists of repetitive elements and that those repetitive elements are more abundant in introns than exons, likely contributing to the evolution of gene sizes. Below we discuss how our results contribute to our understanding of the evolution of genome structure in frogs and highlight an application of the *D. tinctorius* genome to facilitate marker development for cost-effective, population-scale multilocus genotyping using MobiSeq (Rey-Iglesia et al. 2019).

### Transposable elements and genome evolution

Amphibians are particularly useful models to provide insights into relationships between environmental or ecological factors and the dynamics of genome size evolution (Liedtke et al. 2018). Transposable elements are an interesting component of genome architecture, as their abundance and diversity has the potential to contribute to genetic diversity and subsequent adaptations and divergence within and between species (Schrader and Schmitz 2018; Ding et al. 2016). We find that over three quarters of the *D. tinctorius* genome consists of repetitive elements and that different elements can be found within—or overlapping—both exons and introns (tables 1 and 2). This finding is consistent with the few studies that have generated chromosome-level genome assemblies for other amphibians. Notably, analyses of the Mexican axolotl (*Ambystoma mexicanum*; 32 Gb genome) and Tibetan frog (*Nanorana parkeri*; 2 Gb genome) genomes have reported LTR elements as the most abundant class of repetitive elements (Nowoshilow et al. 2018; Sun et a. 2015). By contrast, in the relatively small amphibian genome of the Western clawed frog (*Xenopus tropicalis*; 1.5 Gb) LTR elements are less abundant, while DNA transposons are more abundant (Hellsten et al. 2010). The fact that LTR retroelements are abundant in large amphibian genomes has led to the hypothesis that these elements play a particularly important role in the evolution of “genomic gigantism“ (Sun et al. 2012). However, the mechanism underlying the expansion of LTR elements in large amphibian genomes, or the phenotypic consequences of their proliferation, remain to be tested. Our analyses of LTR abundances, size, and insertion times corroborate past findings and indicated that LTRs have played a significant role in the genome size evolution of *D. tinctorius* (fig 4). The fact that many repetitive elements are within or overlap gene features (introns and exons; tables 1 and 2) suggests that their evolution may have important phenotypic consequences. For example, studies in other non-amphibian species have shown that TEs can have diverse effects on gene expression (Lanciano and Cristofari 2020; Rech et al. 2022); however, more work is needed to understand the phenotypic consequences of TEs across ecologically and behaviourally diverse amphibian species.

SINEs are a highly abundant class of transposable element in mammals and many other vertebrates, although they are almost entirely absent in amphibian genomes (Chalopin et al. 2015; Zuo et al. 2023), including *D. tinctorius*. This difference illustrates how the abundance of different transposable elements varies greatly across the tree of life. Interestingly, the majority of transposable elements we annotated in the *D. tinctorius* genomes could not be classified as known elements using our approach (39.15%). This is likely because reference libraries of described transposable elements used in annotation pipelines lack amphibian-specific elements, and this pattern of abundant “unclassified” elements is common in studies of amphibian genomes (Sotero-Caio et al. 2017; Zuo et al. 2023). Future work that leverages ever-increasing amphibian genomic resources to describe the diversity and structure of the amphibian ‘dark matter’ elements would open doors for comparative analyses of transposable element evolution across taxa and greatly increase our understanding of the evolutionary history of parasitic DNA across the tree of life.

A proximate consequence of TE evolution is their effect on genome size. For example, genome size increases with the abundance of repetitive elements in anurans (Zuo et al. 2023), thus suggesting that recent introduction, and lack of deletion, of these elements has led to an increase in genome sizes. Our results show that effects on genome size are not restricted to intergenic regions, with TEs being found within both exons and introns (tables 1 and 2). We also find that TEs can affect the size of gene regions within a genome (fig 3); this finding is contrary to a lack of relationship between gene size and genome size reported in a recent comparison of 14 anuran genomes (Zuo et al. 2023). A possible explanation for this discrepancy is that the largest genome in Zuo et al. (2023)’s analysis—that of the poison frog *Ranitomeya imitator*— contained SINEs and was considerably larger than the size estimated for that species, suggesting abnormalities in the assembly.

While it is now clear that TEs comprise a large proportion of many amphibian genomes, and they are not restricted to intergenic regions (Sotero-Caio et al. 2017), there is still a knowledge gap in our understanding of how TEs influence the evolution and adaptability in this group. A possible mechanism linking aspects of genome size evolution—such as the accumulation of TEs in introns—and phenotypic evolution, is the effect that intron size can have on gene expression (Castillo-Davies et al. 2002; Taft et al. 2007). Indeed, many TEs present in poison frog genomes are actively expressed (Rogers et al. 2018). Future work that leverages an increasing number of amphibian genomes with transcriptomic analysis—for example analysing expression of different isoforms across species and developmental stages—could provide a fruitful avenue towards addressing the ‘consequences of genome-size evolution’ knowledge gap.

### Multilocus genotyping using MobiSeq

We used our genome assembly to show that leveraging repetitive elements to design and anchor primers may be a useful way to generate multilocus genotypic data at the population scale. These types of datasets would open exciting possibilities for understanding the adaptive processes and evolutionary dynamics of amphibian populations. The low amount of DNA that is needed for sequencing over larger parts of the genome makes MobiSeq a powerful tool for non-invasive sampling of target species. The target primers can be developed from the reference genome (if available), or from closely related species, as we have shown in our study. However, we recommend using a reference genome whenever possible to design species-specific primers and to map reads and genotype SNPs.

Our test of the MobiSeq approach showed that primers can differ considerably in their amplification and sequencing success. The element LINE109 could have been not abundant enough throughout the genome to give enough sequences: the LINE109 element was found 109 times in the *D. tinctorius* assembly, while the unknown transposable element TE644 was found 644 times. Therefore, we suggest that future marker development should design primers for markers which have at least 400 to 600 copies in the genome.

When genotyping and calling SNPs, de novo assembly with the program stacks gives better results than mapping to the *O. pumilio* genome where the primers were developed from (the *Dendrobates tinctorius* assembly was not available when we started the MobiSeq test). Although this is a closely related species, high variability in TE composition could contribute to poorer mapping and genotyping when relying on the *O. pumilio* genome. The de novo approach also gave better results than the mapping to the new *D. tinctorius* reference genome. A limitation of the stacks method we used for genotyping is that this approach resulted in high genotype missingness. Additionally, the stacks program was developed for restriction site digested (RAD) approaches; therefore, we highly recommend using the original ANGSD pipeline, provided by the team that developed MobiSeq when genotyping MobiSeq data (Rey-Iglesias et al. 2019). This method gave the best results considering genotyping and further analysis with COLONY. Specifically, filtering SNPs to those called within a minimum of 70 (out of 92) individuals gave the most reliable relationship estimates in terms of identifying 5/5 of the duplicates in our dataset, as well as giving realistic population-size estimates, given our knowledge of the sample population and the biology of *D. tinctorius*.

MobiSeq was developed using mammalian genomes, which have lower levels of transposable elements, and focused on SINEs and LINEs, which are less common in amphibians. Therefore, there are challenges in applying this method to amphibians with large genomes and high transposable element content, such as primer selection and mapping to an existing reference genome.

## Conclusions

Transposable elements are a major component of amphibian genomes and they play an important role in genome size evolution. By generating and analysing a long-read assembly of the poison frog, *Dendrobates tinctorius*, we have shown that TE evolution impacts genome size, not only through their insertion in intergenic regions, but also within exons and introns. This “ecology” of TEs provides a possible mechanism that links genome size evolution to phenotypic evolution. We also provide an example use of MobiSeq to generate low-input cost-effective population genetic data. This method could be used to study the evolutionary dynamics of amphibian populations, alongside aiding their conservation. Overall, our study adds to the growing body of knowledge on the evolution of amphibian genomes. We hope the data and analyses we report here will be a valuable resource for future studies of amphibian genetics, evolution, behaviour, and conservation.

## Materials and Methods

### Sample collection, DNA extraction, and sequencing for whole genome assembly

We generated a reference genome for *Dendrobates tinctorius* from a single male of the “azureus” morph / population. This individual was captive bred by hobbyists in the United Kingdom, kept under licence of the Home Office at Bangor University, and sacrificed by overdose of Tricaine Methanesulfonate followed by immediate pithing and decapitation. DNA was extracted in four parallel extractions performed at the NERC Environmental Omics Facility (NEOF) at the University of Sheffield using the Macherey-Nagel NucleoBond High Molecular Weight DNA kit (see SI for details). DNA was then cleaned and sheared to an average size of 19 kb before generating four HiFi libraries using SMRTbell template express kit 2.0 (PacBio). Final libraries were size-selected in the size range of 7-50kb and sequenced across 18 SMRT cells (see SI for details).

### Genome assembly, quality control, and synteny

We assembled HiFi reads greater than 10kb in length using HiFiASM (v0.16.1-r375; Cheng et al. 2021, 2022) run with a bloom filter of 39 bits (option -f f39) and “aggressive” purging of haplotigs (option -l 2). To identify and correct assembly errors, we ran our primary assembly through the Inspector pipeline using inspector.py and inspector-correct.py scripts, respectively (Chen et al. 2021). Inspector also provides an estimate of assembly accuracy, reported as a Quality Value (QV = -10log_10_(base-level errors / total assembly length)). After error-correction we identified and removed bacterial or viral contaminants using Kraken2 (v2.1.2 ; Wood et al. 2019) run using the standard Refseq database containing archaea, bacteria, viral, plasmid, human, and UniVec_Core indexes (downloaded 27-5-2021). We estimated assembly completeness using BUSCO (v5.2.2; Simão et al. 2015) run using the tetrapoda_odb10 dataset, which consists of 5,310 single-copy orthologs derived from 38 genomes (created 2021-02-19). We also compared assembly statistics of the *D. tinctorius* genome to the only other Dendrobatid genome currently available on NCBI (*Oophaga pumilio*) and four chromosome-scale assemblies of other Anurans (*Bufo bufo* [NCBI RefSeq ID: GCF_905171765.1], *Rana temporaria* [RefSeq ID: GCF_905171775.1], *Engystomops pustulosus* [GenBank ID: GCA_019512145.1], and *Xenopus tropicalis* [RefSeq ID: GCF_000004195.4]).

### Synteny between D. tinctorius and other Hyloidea genomes

We compared the *D. tinctorius* assembly to chromosome-scale assemblies of *Bufo gargarizans* (Asiatic toad; Bufonidae; GenBank accession GCA_014858855.1), *Eleutherodactylus coqui* (common coquí; Eleutherodactylidae; GCA_019857665.1), and *Engystomops pustulosus* (túngara frog; Leptodactylidae; GCA_019512145.1) using orthology-guided synteny map construction in GENESPACE (v1.3.1; Lovell et al. 2022) and BUSCO (v5.2.2; Simão et al. 2015) annotations of the 5,310 gene tetrapoda_odb10 dataset (created 2021-02-19) as input. BUSCO annotations for *Xenopus tropicalis* (GCA_000004195.4) were used as outgroup sequences in the initial GENESPACE analysis. Because we were interested in identifying large regions of synteny, we constrained our analysis to scaffolds that have been assigned to chromosomes in the chromosome-scale assemblies, and contigs at least 20 Mb long in the *D. tinctorius* assembly. This approach resulted in the number of annotated single-copy BUSCOs per genome being: 4,482, 3,579, 3,337, 3,440, and 4,939 for *B. gargarizans*, *En. pustulosus*, *El. coqui*, *D. tinctorius*, and *X. tropicalis*, respectively. We present summaries of synteny between *D. tinctorius* and the three chromosome-scale assemblies as the proportion of the *D. tinctorius* assembly contained within a synteny block and the average size of synteny blocks between genomes. Syntenic alignments were visualised using GENESPACE’s plot_riparian function (Lovell et al. 2022).

We also tested whether the size of annotated BUSCO genes was positively correlated with genome size using a Spearman’s rank correlation test. We focused this analysis on BUSCO annotations rather than *de novo* annotations generated for each genome because we annotated BUSCO genes using the same pipeline and BUSCO genes are chosen based on orthology across different vertebrate genomes, thereby reducing the likelihood that comparisons between paralogous genes affected our results. Finally, we compared the *B. gargarizans* assembly—as it was the closest in size to that of *D. tinctorius*—to *D. tinctorius* using a non-ortholog based nucleotide alignment with minimap2 (v2.24-r1122; Li 2018, 2021; options:-x asm20 -I10G -B3 -O4,24 -N10).

### Genome annotation and transposable element diversity

We annotated repetitive elements in our *D. tinctorius* assembly by first using RepeatModeler (v2.0.3; Flynn et al. 2020) to identify and generate a species-specific repeat library, followed by RepeatMasker (v 4.1.2-p1; Smith et al. 2021) run using the custom repeat library generated by RepeatModeler. We also conducted an independent annotation of Long Terminal Repeat (LTR) retrotransposons using LtrDetector (Valencia & Girgis 2019). We used BLAST and the custom repeat library generated by RepeatModeler to classify LTR retroelements annotated by LtrDetector. To estimate LTR insertion times we extracted left and right LTR regions from the *D. tinctorius* genome in fasta format using bedtools (Quinlan & Hall 2010) and the annotated coordinates generated by LtrDetector. We retained annotations where BLAST was able to match greater than 50% of the LTR retroelement with an element contained within the library generated by RepeatModeler (v2.0.3; Flynn et al. 2020). We then aligned left and right elements for each LTR retroelement using MAFFT (v7.490; Katoh & Standley 2013; options: --globalpair --maxiterate 1000) and estimated insertion times from divergence between the left and right LTR, which we estimated from each MAFFT alignment using the dist.dna function in R run assuming Kimura’s 2-parameter model (Kimura 1980; option: model = “K80”). To convert divergence estimates to time, we assumed a substitution rate of 2.5 x 10^-9 substitutions per site per year (Lau et al. 2020).

In addition to repeat elements, we annotated protein-coding genes *de novo* using the BRAKER2 pipeline (v2.1.6; Brůna et al. 2021). Before annotation we masked the genome using RepeatModeler and RepeatMasker with default settings. We run BRAKER2 using evidence from 22 RNAseq libraries from brain, eggs, gut, liver, and skin. RNAseq reads were trimmed using Trimmomatic (v. 0.39; Bolger et al. 2014) to remove adaptor contamination and low-quality bases (options: LEADING:9 TRAILING:9 SLIDINGWINDOW:4:15 MINLEN:80). Trimmed reads were then mapped to the genome using STAR (v. 2.7.8a; Dobin et al. 2013) specifying the twopass Mode Basic option. Mapped RNAseq reads were then used as evidence in the BRAKER2 pipeline. UTR predictions were added to these predictions using the --addUTR=on option. Finally, UTR annotations were trimmed following a gap of over 1000bp. We used these annotations—alongside those generated by RepeatModeler + RepeatMasker—to test whether repetitive elements played a role in the evolution of intron / exon structure in the *D. tinctorius* genome using bedtools (Quinlan & Hall 2010).

### Tadpole sampling and DNA extraction for relatedness analysis

The aim of this part of the study was to test the MobiSeq protocol as a method for generating a reduced representation library in a non-model organism, the dyeing poison frog (*Dendrobates tinctorius*). We wanted to use single nucleotide polymorphism (SNP) data to resolve the relatedness between tadpoles in different, small, confined environments such as phytotelmata (small bodies of water in tree holes or palm bracts, for natural history see Rojas and Pašukonis 2019).

We sampled 87 tadpoles of *D. tinctorius* from 17 phytotelmata in the Nouragues Nature Reserve, French Guiana in 2020, by clipping a small part of the tail tip. Tissue samples were stored in 70% ethanol (EtOH) at -20°C until further processing. To increase reliability and to test the appropriateness of the relatedness analysis, we included 5 samples as duplicates, giving a total sample size of 92. A modified salting-out method was used to extract DNA from small parts of the tail clips (supplementary methods, section 1, Supplementary Material online). The extracted DNA was eluted in 100µl TE buffer, DNA concentration measured with a NanoDrop^TM^ spectrophotometer (Thermo Fisher) and stored at -20°C until further processing.

### MobiSeq primer design

We designed specific primers for repeated elements as MobiSeq uses transposable elements (TEs) in the DNA for target enrichment PCRs. No published reference genome for *D. tinctorius* was available when we started to work on the MobiSeq approach. To find highly repetitive TEs, we used the genome of a closely related species, the strawberry poison frog (*Oophaga pumilio*), which was published in 2018 (Rogers et al.; NCBI GenBank: GCA_009801035.1).

We used the free software RepeatMasker (v 4.1.2; Smith et al. 2021) to mask repeated elements and extracted those regions with Samtools (Danecek et al. 2021) to get the 22 last bp of transposable elements as a reversed complement and the number of occurrences. The list was cross checked and annotated with existing libraries of transposable element families provided by Dfam (v3.2, Storer et al. 2021). Based on this list we choose elements with an average abundance of 400-600 times in the genome, as elements with higher abundances might be clustered and not evenly spread over the genome. LINES and SINEs were generally less abundant so that we included only one primer for a lower abundance LINE. We initially chose six possible reverse transposable element primers and tested amplification and multiplexing with *D. tinctorius* DNA. We decided on the following two primers after running a target enrichment PCR, based on amplification success and minimisation of primer dimers (AdapterTEsequence):

D_tinct_Line_109 (5’GTGACTGGAGTTCAGACGTGTGCTCTTCCGATCTCCTATGTTACTATGTTACTATGT’3)

D_tinct_TE_644 (5’GTGACTGGAGTTCAGACGTGTGCTCTTCCGATCTTACTTTTGGCCACCACTGTA’3).

In the first step of the MobiSeq protocol, the *D. tinctorius* DNA was digested using fragmentase. To ensure an even digestion across samples, the DNA was diluted to 10ng/µl. We tested the correct incubation time for the fragmentase enzyme beforehand and decided on 20 min at 37°C for each sample based on gel images (supplementary methods, section 4.1, Supplementary Material online). After fragmentase treatment, a Sera-Mag Speed beads clean-up was used to remove the enzyme mix and buffers (supplementary methods, section 4.2, Supplementary Material online). Clean-up was followed by end-repair of DNA fragments to create blunt ended fragments using T4 polymerase. After incubation, samples were cooled down to 10°C and immediately used for the next step (supplementary methods, section 4.3, Supplementary Material online). For adapter ligation, double stranded modified P5 adapters (Meyer & Kirchner 2010) were added to the end-repaired DNA fragments via a T4 DNA ligase. The modification of the adapter results in single-stranded adapters, which allow the use of universal adapter primers in the next target enrichment PCR step. The adapter ligation master mix (4µl per sample) is added to the end-repaired samples. The samples were cooled down to 10°C after the incubation (supplementary methods, section 4.4, Supplementary Material online). The resulting product was cleaned again, using the same Sera-Mag Speed beads as after DNA fragmentation (supplementary methods, section 4.2, Supplementary Material online) and eluted in 25µl AE buffer and stored at 4°C. We used the primers TE644 and Line109 as described above to enrich the fragments containing transposable elements in a multiplex approach (supplementary methods, section 4.5, Supplementary Material online). The remaining 15µl of PCR were cleaned using the Sera-Mag Speed beads and eluted in 20µl AE buffer. In the final step of the sequencing library preparation, a second PCR using Illumina indexed forward and reverse primers was used followed by agarose gel electrophoresis (supplementary methods, section 4.6, Supplementary Material online). The uniquely tagged samples were mixed, based on the brightness of the smear on the gel. The whole pool of samples was loaded on another gel to cut out fragments of the lengths between 200 and 500bp. The innuPREP DOUBLEpure kit (Analytik Jena) was used to extract the DNA from the gel fragment, following the kit’s protocol. The final library pool was sent to Novogene UK for sequencing in 150bp paired end mode on a Novaseq instrument (Illumina).

### Bioinformatics for variant calling

After downloading the demultiplexed fastq files from Novogene we visually checked the reads for quality with qiime2 (v2021.4; Bolyen et al. 2019) using the demux plugin (function summarize). As each fastq file contains Line109 and TE644 reads, we used cutadapt (version 4.1, Martin 2011) to separate each file into two, based on the primer sequence. After separating the files, we used cutadapt to remove all versions of adapters and primers from the forward and reverse reads. Reverse-complement versions were used to remove primer fragments that can be present on the 5’ end of reads from fragments that are shorter than 150bp. After trimming, the sequences were filtered for optical duplicates and remains of bacteriophage PhiX contamination using the shell scripts clumpify.sh and bbduke.sh from BBTools (Bushnell 2014) .

For reduced representation sequencing approaches such as MobiSeq to be broadly applicable to non-model species that lack genomic resources, it is important to know how reliant markers are on the availability of genomic resources of closely related taxa. As such, we took three approaches to calling SNPs in our dataset: (1) de novo assembly of MobiSeq fragments, (2) mapping reads to the *O. pumilio* reference, and (3) mapping to the *D. tinctorius* genome we generated as part of this study. The de novo assembly was conducted twice using denovo_map.pl from the stacks pipeline (v2.64; Catchen et al. 2011, 2013), using the default parameters. Therefore, all sequences were truncated to 50bp and 100bp, respectively for both primers using process_radtags. The reason for the double truncation is that the truncation process discards sequences shorter than the chosen length (50 or 100bp). The 50bp approach therefore retains more but shorter sequences than the 100bp approach. We wanted to compare different truncation lengths to validate the robustness of de novo assembly. The reference genome mapping was done with ref_map.pl from the stacks pipeline (v2.64; Catchen et al. 2011, 2013), using the default parameters. Before mapping, the reference genomes were indexed with the BWA-mem2 index function (version 2.2.1; Vasimuddin et al. 2019) and subsequently sequences aligned with the mem function. The output was saved as a vcf file, with the following parameters: populations: -p 1 -r 0 --write-random-snp --max-obs-het 0.5 --ordered-export --vcf.

The soft filtering of SNP data was done in R version 4.2.2 (R Core Team 2022) with Bioconducter version 3.16 (BiocManager 1.30.19; Morgan 2022) and the filter_rad function from the radiator package (v1.2.8; Gosselin, 2020). To implement the radiator package from github we used the package devtools (Wickham et al. 2022). All thresholds that we used for filtering can be found in supplementary methods (section 5, Supplementary Material online). We expected a high degree of missing data due to the Mobiseq approach, therefore the filter for maximum missingness was quite high (0.9).

The strong filtering of SNP data was done with vcftools (v. 0.1.16; Danecek et al. 2011) using the following parameters: --max-missing 0.5 --mac 2 --minDP 3.

Additionally, we used the ANGSD pipeline from Rey-Iglesia et al. 2019 (https://github.com/shyamsg/MobiSeq/blob/master/code/pipeline.sh), accounting for the fact that stacks was developed for restriction enzyme based RAD sequencing and might bias our output. We used the BWA-mem2 indexed reference genome of *D. tinctorius* to map our sequences, bedtools (v 2.26.0; Quinlan & Hall 2010) to merge reads and kept only sites that are present in 90% of the cases. We called variants using ANGSD (v 0.940, Korneliussen et al. 2014) with minimum quality of 30, min mapping quality of 30, filter for SNPs with a p-value of 1e-6, major and minor allele were inferred directly from likelihoods, minor allele frequency was estimated (fixed major unknown minor), genotypes and SNPs called. SNPs needed to be present in either a minimum of 80 or 70 individuals. We randomly called one SNP per contig to avoid linkage and created 5 vcf files for the 80 and 70 individuals dataset respectively.

### Relatedness analysis

The relatedness analysis was conducted with COLONY 2.0 (Jones and Wang 2010). We used the filtered SNPs of the TE644 primer only, derived from our 4 approaches (de novo stacks, *O. pumilio* reference genome or our new *D. tinctorius* genome) and two methods of calling SNPs (stacks vs. ANGSD). The write_colony function in the radiator package was used to write a colony input function. The input parameters were as follows; the mating system of males and females was set to polygamous, no inbreeding and no update of allele frequencies, the length of the run was set to 2 with full likelihood analysis and high precision. We ran an analysis with different random seeds to increase reliability of the clustering (1234 and 1789).

## Acknowledgements

We thank Lia Schlippe for providing drawings of *Oophaga pumilio* and *Dendrobates tinctorius*. Silhouettes of different frogs presented in Figures 1, 2 and 3 were taken from the PhyloPic website (phylopic.org). The DNA icon was taken from flaticon.com. This work was supported by funding from the NERC Environmental Omics Facility (project number 1382 to AAC); Research Council of Finland (# 318404; 319949; 345974 to BR); and the National Science Foundation (IOS-1822025 and IOS-1827333 to LAO).

## Data availability

The genome assembly of *D. tinctorius* is publicly available on NCBI: BioProject: PRJNA977854; BioSample: SAMN35537897. Illumina reads generated from MobiSeq libraries are publicly available on NCBI: BioProject: PRJNA1035148.

## Authors’ contributions

AAC, SS and BR conceived and designed the project. RM, CAF and AAC collected samples. CD, FH, LSK, MH extracted DNA and prepared the libraries. SS, LAO, AF, AAC provided resources (data, computational). DP, FH, CD, AAC performed bioinformatic analysis of the data. CD, AAC, FH, BR and SS discussed and interpreted the results, AAC, CD, FH and LAO wrote the first draft. AAC and BR provided funding. All authors read, reviewed and approved the final manuscript.

## References

1. Almojil D et al. 2021. The structural, functional and evolutionary impact of transposable elements in eukaryotes. Genes. 12(6):918.

2. Amézquita A, Flechas SV, Lima AP, Gasser H, Hödl W. 2011. Acoustic interference and recognition space within a complex assemblage of dendrobatid frogs. Proc Natl Acad Sci USA. 108:17058–17063.

3. Bourque G et al. 2018. Ten things you should know about transposable elements. Genome Biol. 19:1–2.

4. Bolger AM, Lohse M, Usadel B. 2014. Trimmomatic: a flexible trimmer for Illumina sequence data. Bioinformatics. 30:2114–2120.

5. Bolyen E et al. 2019. Reproducible, interactive, scalable and extensible microbiome data science using QIIME 2. Nature Biotechnol. 37: 852–857.

6. Brandies P et al. 2019. The value of reference genomes in the conservation of threatened species. Genes. 10(11): 846.

7. Brůna T, Hoff KJ, Lomsadze A, Stanke M, Borodovsky M. 2021. BRAKER2: automatic eukaryotic genome annotation with GeneMark-EP+ and AUGUSTUS supported by a protein database. NAR Genom Bioinform. 3: lqaa108.

8. Bushnell B. 2014. BBMap short read aligner, and other bioinformatic tools. https://sourceforge.net/projects/bbmap/.

9. Carvajal-Castro JD et al. 2021. Aposematism facilitates the diversification of parental care strategies in poison frogs. Sci Rep. 11(1): 19047.

10. Casacuberta E, González J. 2013. The impact of transposable elements in environmental adaptation. Mol Ecol. 22(6):1503–17.

11. Castillo-Davis CI, Mekhedov SL, Hartl DL, Koonin EV, Kondrashov FA. 2002. Selection for short introns in highly expressed genes. Nat Genet. 31:415–418.

12. Catchen J, Amores A, Hohenlohe P, Cresko W, Postlethwait J. 2011. *S*tacks: building and genotyping loci de novo from short-read sequences. G3: Genes, Genomes, Genetics. 1:171–182.

13. Catchen J, Hohenlohe P, Bassham S, Amores A, Cresko W. 2013. Stacks: an analysis tool set for population genomics. Mol Ecol. 22:3124–3140.

14. Chalopin D, Naville M, Plard F, Galiana D, Volff JN. 2015. Comparative analysis of transposable elements highlights mobilome diversity and evolution in vertebrates. Genome Biol Evol. 7(2):567–80.

15. Chen Y, Zhang Y, Wang AY, Gao M, Chong Z. 2021. Accurate long-read de novo assembly evaluation with Inspector. Genome Biol. 22:312.

16. Cheng H, Concepcion GT, Feng X, Zhang H, Li H. 2021. Haplotype-resolved de novo assembly using phased assembly graphs with hifiasm. Nature Methods. 18(2):170–5.

17. Cheng H et al. 2022. Haplotype-resolved assembly of diploid genomes without parental data. Nature Biotechnol. 40(9):1332–1335.

18. Chouteau M, Summers K, Morales V, Angers B. 2011. Advergence in Müllerian mimicry: the case of the poison dart frogs of Northern Peru revisited. Biol Lett. 7(5):796–800.

19. Danecek P et al. 2011. The variant call format and VCFtools. Bioinformatics. 27(15):2156–2158.

20. Danecek P et al. 2021. Twelve years of SAMtools and BCFtools. GigaScience. 10(2), giab008 [33590861].

21. Ding Y, Berrocal A, Morita T, Longden KD, Stern DL. 2016. Natural courtship song variation caused by an intronic retroelement in an ion channel gene. Nature. 536:329–332.

22. Dobin A et al. 2013. STAR: ultrafast universal RNA-seq aligner. Bioinformatics. 29:15–21.

23. Donnelly MA. 1989. Effects of reproductive resource supplementation on space-use patterns in *Dendrobates pumilio*. Oecologia. 81:212–218.

24. Dufresnes C et al. 2018. Genomic evidence for cryptic speciation in tree frogs from the Apennine Peninsula, with description of *Hyla perrini* sp. nov. Front Ecol Evol. 2018:144.

25. Feng YJ et al. 2017. Phylogenomics reveals rapid, simultaneous diversification of three major clades of Gondwanan frogs at the Cretaceous–Paleogene boundary. Proc Natl Acad Sci USA. 114(29):E5864–70.

26. Feschotte, C. 2008. Transposable elements and the evolution of regulatory networks. Nat Rev Genet. 9:397–405.

27. Fischer EK et al. 2019. The neural basis of tadpole transport in poison frogs. Proc Soc B. 286:20191084.

28. Fischer EK, O’Connell LA. 2020. Hormonal and neural correlates of care in active versus observing poison frog parents. Horm Behav. 120:104696.

29. Fischer EK et al. 2020. Neural correlates of winning and losing fights in poison frog tadpoles. Physiol Behav. 223:112973.

30. Flynn JM et al. (2020). RepeatModeler2 for automated genomic discovery of transposable element families. Proc Natl Acad Sci USA. 117: 9451–9457.

31. Fouilloux C et al. 2021. Pool choice in a vertical landscape: tadpole rearing site flexibility in phytotelm-breeding frogs. Ecol Evol. 11: 9021–9038.

32. Gozashti L, Feschotte C, Hoekstra HE. 2023. Transposable Element Interactions Shape the Ecology of the Deer Mouse Genome. Mol Biol Evol. 40:msad069.

33. Gosselin T. 2023. radiator: RADseq Data Exploration, Manipulation and Visualization using R. R package version 1.2.8 https://thierrygosselin.github.io/radiator/

34. Hawkins JS, Kim H, Nason JD, Wing RA, Wendel JF. 2006. Differential lineage-specific amplification of transposable elements is responsible for genome size variation in Gossypium. Genome Res. 16(10):1252–61.

35. Hellsten U. et al. 2010. The genome of the Western clawed frog *Xenopus tropicalis*. Science. 328(5978):633–636.

36. Homola JJ et al. 2019. Replicated Landscape Genomics Identifies Evidence of Local Adaptation to Urbanization in Wood Frogs. J Heredity. 110:707–719.

37. Hotaling S, Kelley JL, Frandsen PB. 2021. Toward a genome sequence for every animal: where are we now?. Proc Natl Acad Sci USA. 118(52):e2109019118.

38. Jones O, Wang J. 2010. COLONY: a program for parentage and sibship inference from multilocus genotype data. Mol Ecol Res. 10:551–555.

39. Katoh K, Standley DM. 2013. MAFFT multiple sequence alignment software version 7: improvements in performance and usability. Mol Biol Evol. 30:772–780

40. Kidwell MG, Lisch D. 1997.Transposable elements as sources of variation in animals and plants. Proc Natl Acad Sci USA. 94(15):7704–7711.

41. Kidwell MG. 2002. Transposable elements and the evolution of genome size in eukaryotes. Genetica. 115:49–63.

42. Kimura M. 1980. A simple method for estimating evolutionary rates of base substitutions through comparative studies of nucleotide sequences. J Mol Evol, 16:111–120.

43. Korneliussen TS, Albrechtsen A, Nielsen R. 2014. ANGSD: Analysis of Next Generation Sequencing Data. BMC Bioinformatics. 15:356.

44. Kosch TA et al. 2022. Genetic approaches for increasing fitness in endangered species. Trends Ecol Evol. 37:332–345.

45. Lamichhaney S et al. 2021. A bird-like genome from a frog: Mechanisms of genome size reduction in the ornate burrowing frog, *Platyplectrum ornatum*. Proc Natl Acad Sci USA. 118(11):e2011649118.

46. Lanciano S, Cristofari G. 2020. Measuring and interpreting transposable element expression. Nat Rev Genet. 21:721–736.

47. Lau Q et al. 2020. Heterogeneity of synonymous substitution rates in the Xenopus frog genome. PLoS One. 15(8):e0236515.

48. Lawrence JP, Rojas B et al. 2019. Weak warning signals can persist in the absence of gene flow. Proc Natl Acad Sci USA. 116(38):19037–45.

49. Lee SI, Kim NS. 2014. Transposable elements and genome size variations in plants. Genomics & Informatics. 12(3):87.

50. Li H. 2018. Minimap2: pairwise alignment for nucleotide sequences. Bioinformatics. 34(18):3094–3100.

51. Li H. 2021. New strategies to improve minimap2 alignment accuracy. Bioinformatics. 37:4572– 4574.

52. Liedtke HC, Gower DJ, Wilkinson M, Gomez-Mestre I. 2018. Macroevolutionary shift in the size of amphibian genomes and the role of life history and climate. Nature Ecol Evol. 2(11):1792– 1799.

53. Lovell JT et al. 2022. GENESPACE tracks regions of interest and gene copy number variation across multiple genomes. eLife. 11:e78526.

54. Maan ME, Cummings ME. 2012. Poison frog colors are honest signals of toxicity, particularly for bird predators. Am Nat. 179(1):E1–4.

55. Martin M. 2011. Cutadapt removes adapter sequences from high-throughput sequencing reads. EMBnet.journal. 17(1):10–12.

56. McClintock B. 1950. The origin and behavior of mutable loci in maize. Proc Natl Acad Sci USA. 36(6):344–355.

57. Meyer M, Kircher M. 2010. Illumina sequencing library preparation for highly multiplexed target capture and sequencing. Cold Spring Harbor Protocols. 2010(6):pdb-rot5448.

58. Morgan M. 2020. BiocManager: Access the Bioconductor Project Package Repository. R package version 1.30.19, https://CRAN.R-project.org/package=BiocManager

59. Myers CW, Daly JW. 1983. Dart-poison frogs. Sci Am. 248:96–105.

60. Noonan BP, Comeault AA. 2009. The role of predator selection on polymorphic aposematic poison frogs. Biol Lett. 5(1):51–4.

61. Nowoshilow S, et al. 2018. The axolotl genome and the evolution of key tissue formation regulators. Nature. 554:50–55.

62. Nunziata SO, Weisrock DW. 2018. Estimation of contemporary effective population size and population declines using RAD sequence data. Heredity. 120(3):196–207.

63. Pašukonis A et al. 2022. Contrasting parental roles shape sex differences in poison frog space use but not navigational performance. eLife. 11:e80483.

64. Pimpinelli, S, Piacentini, L. 2020. Environmental change and the evolution of genomes: Transposable elements as translators of phenotypic plasticity into genotypic variability. Funct Ecol. 34: 428–441.

65. Platt RN, Vandewege MW, Ray DA. 2018. Mammalian transposable elements and their impacts on genome evolution. Chromosome Res. 26:25–43.

66. Pröhl H. 2005. Clutch loss affects the operational sex ratio in the strawberry poison frog *Dendrobates pumilio*. Behav Ecol Sociobiol. 58:310–315.

67. Quinlan AR, Hall IM. 2010. BEDTools: a flexible suite of utilities for comparing genomic features. Bioinformatics. 26:841–842.

68. R Core Team. R: A language and environment for statistical computing. R Foundation for Statistical Computing, Vienna, Austria. 2020. URL https://www.R-project.org/. Version 4.2.2

69. Rech GE et al. 2022. Population-scale long-read sequencing uncovers transposable elements associated with gene expression variation and adaptive signatures in Drosophila. Nature Commun. 13(1):1948.

70. Rey O, Danchin E, Mirouze M, Loot C, Blanchet S. 2016. Adaptation to global change: a transposable element–epigenetics perspective. Trends Ecol Evol. 31(7):514–526.

71. Rey-Iglesia A et al. 2019. MobiSeq: De novo SNP discovery in model and non[model species through sequencing the flanking region of transposable elements. Mol Ecol Res. 19(2):512–25.

72. Ringler E, Pašukonis A, Hödl W, Ringler M. 2013. Tadpole transport logistics in a Neotropical poison frog: indications for strategic planning and adaptive plasticity in anuran parental care. Front Zool. 10:67.

73. Rodríguez C et al. 2022. Androgen responsiveness to simulated territorial intrusions in *Allobates femoralis* males: evidence supporting the challenge hypothesis in a territorial frog. Gen Comp Endocrinol. 326:114046.

74. Rojas B. 2014. Strange parental decisions: fathers of the dyeing poison frog deposit their tadpoles in pools occupied by large cannibals. Behav Ecol Sociobiol. 68:551–559.

75. Rojas B. 2015. Mind the gap: treefalls as drivers of parental tradeoffs. Ecol Evol. 5:4028–4036.

76. Rojas B & Pašukonis A. 2019. From habitat use to social behavior: natural history of a voiceless poison frog, *Dendrobates tinctorius*. PeerJ. 7:e7648.

77. Rogers RL et al. 2018. Genomic takeover by transposable elements in the strawberry poison frog. Mol Biol Evol. 35:2913–2927.

78. Schloissnig S et al. 2021. The giant axolotl genome uncovers the evolution, scaling, and transcriptional control of complex gene loci. Proc Natl Acad Sci USA. 118(15):e2017176118.

79. Schrader L, Schmitz J. 2019. The impact of transposable elements in adaptive evolution. Mol Ecol. 28(6):1537–49.

80. Schulte LM, Lötters S. 2013. The power of the seasons: rainfall triggers parental care in poison frogs. Evol Ecol. 27:711–723.

81. Serrato-Capuchina A, Matute DR. 2018. The Role of Transposable Elements in Speciation. Genes. 9(5):254.

82. Sexton OJ. 1960. Some aspects of the behavior and of the territory of a dendrobatid frog, *Prostherapis trinitatis*. Ecology. 41:107–115.

83. Silverstone PA. 1973. Observations on the behavior and ecology of a Colombian poison-arrow frog, Kõkoé-pá (*Dendrobates histrionicus* Berthold). Herpetologica. 29:295–301.

84. Simão FA, Waterhouse RM, Ioannidis P, Kriventseva EV, Zdobnov EM. 2015. BUSCO: assessing genome assembly and annotation completeness with single-copy orthologs. Bioinformatics. 31(19):3210–3212.

85. Smith A, Hubley R, Green P. RepeatMasker version 4.1.2-p1. 2021. http://repeatmasker.org

86. Sotero-Caio CG, Platt RN, Suh A, Ray DA. 2017. Evolution and diversity of transposable elements in vertebrate genomes. Genome Biol Evol. 9(1):161–177.

87. Stitzer MC, Anderson SN, Springer NM, Ross-Ibarra J. 2021. The genomic ecosystem of transposable elements in maize. PLoS Genet. 17(10):e1009768.

88. Storer J, Hubley R, Rosen J, Wheeler TJ, Smit AF. 2021. The Dfam community resource of transposable element families, sequence models, and genome annotations. Mobile DNA. 12:2.

89. Stynoski JL, Torres-Mendoza Y, Sasa-Marin M, Saporito RA. 2014. Evidence of maternal provisioning of alkaloid[based chemical defenses in the strawberry poison frog *Oophaga pumilio*. Ecology. 95(3):587–93.

90. Stynoski JL, Schulte LM, Rojas B. 2015. Poison frogs. Quick Guide. Curr Biol. 25:R1026– R1028.

91. Summers K. 1989. Sexual selection and intra-female competition in the green poison-dart frog, *Dendrobates auratus*. Anim Behav. 37:797–805.

92. Summers K, Sea McKeon C, Heying H. 2006. The evolution of parental care and egg size: a comparative analysis in frogs. Proc R Soc B. 273:687–692.

93. Sun C et al. 2012. LTR retrotransposons contribute to genomic gigantism in plethodontid salamanders. Genome Biol Evol. 4(2):168–183.

94. Sun YB et al. 2015. Whole-genome sequence of the Tibetan frog *Nanorana parkeri* and the comparative evolution of tetrapod genomes. Proc Natl Acad Sci USA. 112(11):E1257–1262.

95. Taft RJ, Pheasant M, Mattick JS. 2007. The relationship between non[protein[coding DNA and eukaryotic complexity. Bioessays. 29(3):288–99.

96. Tumulty J, Morales V, Summers K. 2014. The biparental care hypothesis for the evolution of monogamy: experimental evidence in an amphibian. Behav Ecol. 25(2):262–70.

97. Valencia JD, Girgis HZ. 2019. LtrDetector: A tool-suite for detecting long terminal repeat retrotransposons de-novo. BMC Genomics. 20:450.

98. Vasimuddin M, Sanchit M, Heng L, Srinivas A. 2019. Efficient Architecture-Aware Acceleration of BWA-MEM for Multicore Systems. IEEE Parallel and Distributed Processing Symposium (IPDPS). doi:10.1109/IPDPS.2019.00041.

99. Wells KD.1980. Behavioral ecology and social organization of a dendrobatid frog (*Colostethus inguinalis*). Behav Ecol Sociobiol. 6:199–209.

100. Wicker T et al. 2007. A unified classification system for eukaryotic transposable elements. Nature Rev Gen. 8(12):973–82.

101. Wickham H, Hester J, Chang W, Bryan J. 2022. devtools: Tools to Make Developing R Packages Easier. R package version 2.4.5, https://CRAN.R-project.org/package=devtools.

102. Womack MC, Metz MJ, Hoke KL. 2019. Larger genomes linked to slower development and loss of late-developing traits. Am Nat. 194(6):854–64.

103. Wood DE, Lu J, Langmead B. 2019. Improved metagenomic analysis with Kraken 2. Genome Biol. 20:1–3.

104. Zuo B, Nneji LM, Sun YB. 2023. Comparative genomics reveals insights into anuran genome size evolution. BMC Genomics. 24(1):379.

