## Supplementary methods for "Genome assembly of the dyeing poison frog provides insights into the dynamics of transposable element and genome-size evolution"

**Content**

1. DNA extraction for the reference genome of *Dendrobates tinctorius*	1
2. Library preparation for sequencing of the reference genome	2

```
3. Protocol to extract DNA from tadpole tissues 		3
4. MobiSeq protocol									4
4.0 Used Oligos, Primers, buffers				4
4.1 Fragmentation								9
4.2 Magnetic bead clean up				   10
4.3 End repair							   11
4.4 Adapter ligation						   12
4.5 Transposable element primer PCR		   13
4.6 Illumina indexing PCR 				   14
5. SNP filtering options with filter_rad			   15
6. Supplementary Tables(captions)				   16
```

1. **DNA extraction for the reference genome of *Dendrobates tinctorius*:**

Muscle tissue from the leg was ground in liquid nitrogen and—to reduce any HMW DNA degradation—immediately placed into lysis buffer. Both lysis buffer and proteinase K volume was doubled, compared to kit’s protocol. The sample was incubated for one hour with gentle agitation every 15 minutes. Binding buffer volume was also doubled and the rest of the process completed per kit instructions. Each extraction was then cleaned using AMpure PB beads and HMW DNA quality was assessed using an Agilent Femto Pulse which showed the majority of DNA greater than 50,000bp in length. Across all four extractions, we obtained a total of 56.9mg of DNA (Qubit Fluorometer), with a purity score of 1.79 (absorbance at 260/280 nm, NanoDropTM spectrophotometer (both Thermo Fisher)).

2. **Library preparation for sequencing of the reference genome:**

DNA was checked for integrity and quality using Femto Pulse, Qubit, and Nanodrop (Thermo Fisher). The Femto Pulse was used throughout the protocol for size checks and Qubit for concentration measurements. For each library, 5µg of DNA in EB buffer (Qiagen) was sheared using the Megaruptor 3 (Diagenode), at a speed code setting of 29 in 500µl volume. The average fragment size was checked and found to be 19kb. This sample was Ampure bead cleaned and used as the starting material for the library preparation, following the manufacturer’s directions. Essentially, this involved a single strand removal step at 37°C for 15 min, DNA damage repair incubation at 37°C for 30 min and an end repair step at 20°C for 30 min and 65°C for 30 min. The SMRT bell overhang adapter was ligated in a reaction at 20°C for 1 h followed by a termination step at 65°C for 10 min. A cocktail of nucleases was added to this reaction and incubated at 37°C for 1 h to remove any non-circular DNA. The sample was Ampure bead cleaned and resuspended in 30µl EB buffer. The sample was size-selected using the Sage blue pippin system, using a 0.75% cassette and a size range 7-50kb in an improved recovery mode. The recovered material had an average size of 18kb.

The size selected library was sequenced on the Pacific Biosciences Sequel 2e. To make the library sequence compatible, a binding complex was made using the Sequel binding kit 2.2, following the guidelines from the SMRT link software. Sample was primed with V5 primer and a complex formed with the Sequel polymerase 2.2. This was ampure bead cleaned and the concentration determined. 50pM on plate loading was used with 30 hour movie times, with 8M SMRT cells. A total of 18 SMRT cells were run with up to 4 libraries, all prepared from the same DNA sample.

**3. Protocol to extract DNA from tadpole tissue**

**DNA extraction of tissue**

\# DNA extraction is a modified salting out method

\# Tissue is stored in EtOH at -20 °C.

\# Extraction negatives need to be included in each round

TNES buffer (50ml):

**NaCl [5M]	4000 µl**	end concentration (400 mM)

**TrisHCl[3M] pH 7.5	833 µl**	end concentration (50 mM)

**SDS [20%]	1250 µl**	end concentration (0.5%)

**EDTA [0.5M]	2000 µl**	end concentration (20 mM)

**Top up** with milliQ water **to 50 ml**

Procedure:

1 Cut off a tiny piece of tissue and dry it to remove the EtOH.
2 Add 500 µl TNES buffer and 20 µl Prot K to the tissue.
3 Vortex and incubate for 20 min at 55°C while shaking (600rpm). The sample should dissolve completely! If not, incubate longer!
4 Spin down quickly
5 add 180 µl 5M NaCl, vortex and spin for 20 min at 12000 rpm (room temp)
6 Put 500 µl of the clear supernatant to a new 1.5 ml tube (containing 1 µl Glycogen and 500 µl chilled isopropanol). **Do not disturb the protein pellet (So take less if 500 µl is not possible)**.
7 Vortex and incubate at -20°C for 5-10 min.
8 Spin for 15 min at 12000 rpm (4°C)
9 Discard supernatant by inverting the tube (do not lose the pellet) and let it sit upside-down on a clean paper towel until all samples are processed.
10 Put the tubes back to the rack and add 500 µl chilled EtOH (70%).
11 Spin for 5 min at 12000 rpm (4°C)
12 Discard supernatant by inverting the tube and let the tube sit upside down on a fresh clean paper towel.
13 Dry the pellet for ~1-2 min in the vacuum centrifuge at 30°C (EtOH must be removed completely). Smell if there is any EtOH left (it is not necessary that all the liquid is dried up as long as it does not smell like EtOH anymore!!!)
14 Dissolve the pellet in 100 µl TE buffer

**4.	MobiSeq protocol**

- This protocol is adapted from Rey‐Iglesia et al. 2019
- all mastermix preparations are carried out on ice to avoid unwanted enzyme activity
- all recipes are shown for one single sample

Used DNA-oligos:

*Oligos to create double stranded mP5 adapter*

**IS1** (Meyer and Kircher (2010))

A*C*A*C*TCTTTCCCTACACGACGCTCTTCCG*A*T*C*T

**modified IS3** (Meyer and Kircher (2010), modified by Rey‐Iglesia et al 2019)

A*G*A*T*CGGAA*G*A*G*C*[SpC3]

(* indicate oligos with phosphorothioate bonds (PTOs); [SpC3] indicate a 3’ Spacer C3)

*Transposable element (TE) primers used in this study*

**D_tinct_Line_109**

GTGACTGGAGTTCAGACGTGTGCTCTTCCGATCTCCTATGTTACTATGTTACTATGT

**D_tinct_TE_644**

GTGACTGGAGTTCAGACGTGTGCTCTTCCGATCTTACTTTTGGCCACCACTGTA

**IS4** (Meyer and Kircher (2010))

AATGATACGGCGACCACCGAGATCTACACTCTTTCCCTACACGACGCTCTT

**P5 indexing primer (1-8)**

P5_ind_1

AATGATACGGCGACCACCGAGATCTACACAATGCCAACACTCTTTCCCTACACGACGCTCT

P5_ind_2

AATGATACGGCGACCACCGAGATCTACACCGGTATCACACTCTTTCCCTACACGACGCTCT

P5_ind_3

AATGATACGGCGACCACCGAGATCTACACTCAGTAGACACTCTTTCCCTACACGACGCTCT

P5_ind_4

AATGATACGGCGACCACCGAGATCTACACCCCAAATACACTCTTTCCCTACACGACGCTCT

P5_ind_5

AATGATACGGCGACCACCGAGATCTACACGGAATCAACACTCTTTCCCTACACGACGCTCT

P5_ind_6

AATGATACGGCGACCACCGAGATCTACACTTCGAGCACACTCTTTCCCTACACGACGCTCT

P5_ind_7

AATGATACGGCGACCACCGAGATCTACACATCGACGACACTCTTTCCCTACACGACGCTCT

P5_ind_8

AATGATACGGCGACCACCGAGATCTACACGACATCTACACTCTTTCCCTACACGACGCTCT

**P7 indexing primer (1-12)**

P7_Ind1

CAAGCAGAAGACGGCATACGAGATCCTGCGAGTGACTGGAGTTCAGACGTGT

P7_Ind2

CAAGCAGAAGACGGCATACGAGATTGCAGAGGTGACTGGAGTTCAGACGTGT

P7_Ind3

CAAGCAGAAGACGGCATACGAGATACCTAGGGTGACTGGAGTTCAGACGTGT

P7_Ind4

CAAGCAGAAGACGGCATACGAGATTTGATCCGTGACTGGAGTTCAGACGTGT

P7_Ind5

CAAGCAGAAGACGGCATACGAGATATCTTGCGTGACTGGAGTTCAGACGTGT

P7_Ind6

CAAGCAGAAGACGGCATACGAGATTCTCCATGTGACTGGAGTTCAGACGTGT

P7_Ind7

CAAGCAGAAGACGGCATACGAGATCATCGAGGTGACTGGAGTTCAGACGTGT

P7_Ind8

CAAGCAGAAGACGGCATACGAGATTTCGAGCGTGACTGGAGTTCAGACGTGT

P7_Ind9

CAAGCAGAAGACGGCATACGAGATAGTTGGTGTGACTGGAGTTCAGACGTGT

P7_Ind10

CAAGCAGAAGACGGCATACGAGATGTACCGGGTGACTGGAGTTCAGACGTGT

P7_Ind11

CAAGCAGAAGACGGCATACGAGATCGGAGTTGTGACTGGAGTTCAGACGTGT

P7_Ind12

CAAGCAGAAGACGGCATACGAGATACTTCAAGTGACTGGAGTTCAGACGTGT

Used enzymes (+ manufacturer name and catalog #):

- dsDNA Fragmentase (NEB; Cat #: M0348S)
- Sera-Mag Speed beads (Cytiva; Cat #:65152105050250)
- T4 DNA polymerase (NEB; Cat #: M0203S)
- T4 PNK (Polynucleotide Kinase) (NEB; Cat #: M0201S)
- T4 DNA ligase (NEB; Cat #: M0202S)
- AllTaq Master Mix (Qiagen; Cat #: 203144)

Buffers and reagents to prepare:

**Reaction enhancer**

PEG 4000 [100%]	0.25 g

BSA [50mg/ml]	40.00 μl

NaCl [5M]	80.00 μl

H_2_O	top up to 1 ml

->store at -20°C

**Oligo hybridization buffer (10x)**

EDTA pH8 [100 mM]	10 μl

Tris HCl pH8 [1M]	10 μl

NaCl [5M]	100 μl

H_2_0	880 μl

-> store at -20°C

**CREATION OF DOUBLE STRANDED MODIFIED P5 ADAPTER (mP5)**

mP5 adapter Mix [40 μM] stock solution

IS1 [100 μM]	80 μl

Modified IS3 [100 μM]	80 μl

Oligo hybridization buffer (10x)	20 μl

H_2_O	20 μl

```
200 µL total volume was aliquoted in 4x 50 µL PCR tubes.

Incubation conditions
95°C		 2min
92°C		20 sec
89°C		20 sec
86°C		20 sec
.
.
.
14°C		20 sec
11°C		20 sec
 8°C----infinite hold

NOTE: ramp rate for all steps is set to 0.1°C/sec
```

Working solution of mP5 [25μM]

mP5 adapter Mix [40μM]	50 μl

H_2_O	30 μl

______________________________________________________________

```
1)DNA-fragmentation using Fragmentase enzyme mix (NEB M0348)

Fragmentase master mix
					            1x
```

Fragmentase Buffer	2 μl

dsDNA Fragmentase	2 μl

DNA (**diluted to 10 ng/μl**)	16 μl

Note: we used water to dilute

Incubation conditions

20 min	37°C

Note: No heated lid

After incubation the product is immediately transferred on ice and **2.5 µL [0,5M] EDTA** are added to the samples to stop the reaction (mix (vortex) shortly).

To check for the right fragment size, 5 µL of the product can be loaded to an agarose gel. Fragmentation depends on the species and on quality and quantity of the DNA. Therefore, the incubation time needs to be tested and adjusted for each project!

The desired fragment length is between 150 - 500 bp

**2)Magnetic bead clean-up**

Magnetic bead solution (MBS) [10ml]

100 μl	Sera-Mag Speed beads (washed 2x in 500 μl 1x TE
buffer)

```
1.8 g	PEG-8000
2.0 ml	NaCl [5M]
```

100 μl	TrisHCl [1M] (pH 8)

100 μl	EDTA [100mM]

5 μl	Tween20

H_2_O -> top-up to 10 ml

Note: make sure that all PEG-8000 is dissolved! The magnetic bead solution is stable for several weeks at 4°C.

Workflow:

- Homogenize MBS by vortexing (avoid foam formation)and allow it to adjust to room temperature (RT) for 30 min
- Add 2 sample-volumes of MBS to the samples (for a sample volume of 10 μl add 20 μl MBS)
- Vortex, spin down (shortly!-magnetic beads must be still in solution) and incubate at RT for 5 min
- Place on magnet and wait until the liquid is clear and the beads are attached to the tube wall
- Discard supernatant (while the samples are still on the magnet)
- Leave samples on the magnet and add 150 μl EtOH [70%] and pipet up and down 3 times
- Discard supernatant and repeat the wash step (the samples can be left on the magnet the whole time)
- Discard supernatant and make sure that there is no EtOH remaining
- Air-dry the samples for 5-10 minutes (don’t overdry), or use a Vacuum centrifuge (1 min at 30°C)
- Add 17 μl TE (1x) or AE-buffer [Qiagen] and mix by vortexing
- Incubate overnight at 4°C (this step can be shortened to 30 min)
- Vortex quickly and put back on the magnet
- Collect 16 μl of the clear eluted sample and place in a new tube

```
3)End repair of the DNA samples

Cleaned samples of the last step are the starting material for this part of the protocol

End-repair master mix
									  1x
```

T4 DNA polymerase [3U/μl]	0.2 μl

T4 PNK (Polynucleotide Kinase) [10U/μl]	0.5 μl

dNTP’s [25mM]	0.2 μl

T4 DNA ligase buffer [10x]	2.0 μl

Reaction enhancer	1.1 μl

DNA-template	16.0 μl

Note: the recipe for the reaction enhancer can be found in the beginning of the Mobiseq protocol

Incubation conditions

30 min	20°C

30 min	65°C

infinite hold	4-12°C

**4)Adapter ligation**

The product of the last step is the starting material for this part of the protocol

**!!!Add 1 μl of the double stranded mP5 adapters to the end repaired samples and mix well!!!**

Notes:

- Protocol for creating the double stranded mP5 adapter is found in the beginning of the MobiSeq protocol
- with low amount of starting material -> less adapter can be added (insert:adapter ratio equals 1:20)

```
Adapter ligation master mix
								  1x
```

T4 DNA ligase buffer [10x]	0.5 μl

PEG-4000 [50%]	3.0 μl

T4 DNA ligase [400U/μl]	0.5 μl

**Template & mP5 adapter**	21.0 μl

Incubation conditions

30 min	20°C

10 min	65°C

infinite hold	4-12°C (10°C in our case)

**Magnetic bead clean-up**

The protocol is identical to section 2, but DNA was eluted in 25 μl TE buffer.

Note: the cleaned, adapter ligated product is stable for several days at 4°C or weeks at -20°C

```
5)PCR using the Transposable element primers (TE-PCR)
Adapter ligated and cleaned up samples are the starting material for this step of the protocol.

PCR master mix
								 1x
```

AllTaq Master Mix [4x]	5.0 μl

TE-primer [10μ]	0.5 μl

IS4 (Primer) [10μ]	0.5 μl

DMSO [100%]	0.3 μl

H_2_0	8.7 μl

Cleaned up template	5.0 μl

Note: if multiplexing of the TE-PCR is desired, other TE primers can be added (0.5 μl), and the H_2_0 is reduced by the same amount.

```
Incubation conditions
 2 min		   95°C
```

5 sec	95°C

```
15 sec		   55°C	   25x
10 sec		   72°C
infinite hold	 4-12°C
```

5 μl of the PCR product is checked on an agarose gel and the amount of template DNA can be adjusted if PCR needs to be repeated. The goal of this PCR is a smear in the desired size range.

Note: A low number of cycles in the PCR machine is preferred to amplify many different fragments instead of many copies of the same fragments.

**Magnetic bead clean-up**

The protocol is identical to section 2, but DNA was eluted in 20 μl TE buffer.

```
6)Indexing PCR and sequencing pool preparation

PCR master mix (full reaction)
								 1x
```

AllTaq mix [4x]	5.0 μl

P5 indexing primer (1-8)	0.5 μl

P7 indexing primer (1-12)	0.5 μl

H_2_0	7.0 μl

Cleaned up TE-PCR product	7.0 μl

Notes:

- based on the gel picture the amount of TE-PCR product can be adjusted individually to get roughly the same quantity of fragments per sample. The H_2_0 needs to be adjusted too!
- use the same polymerase as for the TE-PCR!

Incubation conditions

2 min	95°C

```
 5 sec		   95°C
15 sec		   55°C		12x
10 sec		   72°C
infinite hold	 4-12°C
```

5 μl of the indexing PCR product is loaded on an agarose gel and the remaining products of the indexing PCR are pooled according to the brightness of the smear on the gel (less product of bright samples, more product of weak samples).

```
The whole sample pool is loaded to an agarose gel (with one huge well). It is possible to concentrate the pool in a vacuum centrifuge.

Fragments with a length between 200 and 500 bp are cut out and extracted from the gel using the innuPREP DOUBLEpure kit (Analytik Jena), following the manufacturer's instructions.

After controlling quality and quantity with standard library checks (qBit, qPCR,TapeStation) the library pool is ready to be sequenced on an Illumina platform.

5. SNP filter options in the interactive filter_rad function (radiator package in R, Gosslin 2020)

settings as follows
# thresholds from default outlier values for the first filters
# maf 0.05 or mac 2
# total coverage set to min 3 max 30
# filter genotyping, max missingness set to 0.9
# no filter snp position on the read
# no filter on max SNP per locus (its one anyway)
# no short or long distance filtering
# no exclusion of individuals based on heterozygosity
# no duplicated genomes (if we look at relatedness its possible that individuals are closer to each other)
# no hwe filters

6. Supplementary table legends
Table S1
Output of the log files from the stacks v 2.64 method for all four approaches of alignment. Mapping to the genome of Oophaga pumilio (which was used to design the TE primers), the de novo alignment/stacking of 50bp and 100bp respectively, and the mapping to the new reference genome of Dendrobates tinctorius.

Table S2
Comparison of numbers of called single nucleotide polymorphisms (SNPs); stacking de novo with different length (50 and 100bp), to the reference genome of Oophaga pumilio and to the new reference genome of Dendrobates tinctorius. The number of filtered SNPs gives numbers after filtering with either minor allele count of 2 (mac) or minor allele frequency of 0.05 (maf). Additionally, we show partial output from the COLONY runs with the filtered SNP data including the number of clusters (numbers of clusters with probability above 0.8 in brackets), number of full sib families and range of family members, the proportion of duplicates identified, estimates of population size assuming random (population in Hardy-Weinberg equilibrium) and non-random mating (alpha value estimated from genotype data). NA (not available, either due to low amount of marker loci or low variability).
The ANGSD values are given as mean ± sd for 5 randomly called vcf files per approach (minimum of 70 or 80 individuals containing respective SNP sites).
```
